# Rapid shifts in mitochondrial tRNA import in a plant lineage with extensive mitochondrial tRNA gene loss

**DOI:** 10.1101/2021.07.12.451983

**Authors:** Jessica M. Warren, Thalia Salinas-Giegé, Deborah A. Triant, Douglas R. Taylor, Laurence Drouard, Daniel B. Sloan

## Abstract

In most eukaryotes, transfer RNAs (tRNAs) are one of the very few classes of genes remaining in the mitochondrial genome, but some mitochondria have lost these vestiges of their prokaryotic ancestry. Sequencing of mitogenomes from the flowering plant genus *Silene* previously revealed a large range in tRNA gene content, suggesting rapid and ongoing gene loss/replacement. Here, we use this system to test longstanding hypotheses about how mitochondrial tRNA genes are replaced by importing nuclear-encoded tRNAs. We traced the evolutionary history of these gene loss events by sequencing mitochondrial genomes from key outgroups (*Agrostemma githago* and *Silene* [=*Lychnis*] *chalcedonica*). We then performed the first global sequencing of purified plant mitochondrial tRNA populations to characterize the expression of mitochondrial-encoded tRNAs and the identity of imported nuclear-encoded tRNAs. We also confirmed the utility of high-throughput sequencing methods for the detection of tRNA import by sequencing mitochondrial tRNA populations in a species (*Solanum tuberosum*) with known tRNA trafficking patterns. Mitochondrial tRNA sequencing in *Silene* revealed substantial shifts in the abundance of some nuclear-encoded tRNAs in conjunction with their recent history of mt-tRNA gene loss and surprising cases where tRNAs with anticodons still encoded in the mitochondrial genome also appeared to be imported. These data suggest that nuclear-encoded counterparts are likely replacing mitochondrial tRNAs even in systems with recent mitochondrial tRNA gene loss, and the redundant import of a nuclear-encoded tRNA may provide a mechanism for functional replacement between translation systems separated by billions of years of evolutionary divergence.

## Introduction

The existence of multiple genomes within eukaryotic cells necessitates multiple gene expression systems. Protein synthesis occurs separately in endosymbiotically-derived organelles (mitochondria and plastids) and the cytosol. The coding capacity of mitochondrial genomes (mitogenomes) has dwindled from an estimated thousands of genes in the mitochondrial progenitor to only a few dozen in most eukaryotes (Gray 2012). Import of gene products encoded in the nuclear genome compensates for many of these organellar gene losses (Huynen, et al. 2013). Yet despite their reduced gene content, almost all mitogenomes still encode at least some components of the translational machinery (Sloan, et al. 2018).

Clear patterns have emerged in the retention and loss of certain genes involved in organellar translation systems. The genes encoding the large and small subunit mitochondrial ribosomal RNAs (rRNAs) are almost universally retained in the mitogenome, whereas other gene products, such as aminoacyl tRNA synthetases (aaRSs), are exclusively encoded in the nuclear genome and must be imported into the mitochondrial matrix (Sissler, et al. 2005). The genes for transfer RNAs (tRNAs) exhibit much more heterogeneity with respect to their retention in mitogenomes.

A set of 22 mt-tRNA genes that is sufficient to decode all codons has been retained for over 500 million years in some animal mitogenomes (Anderson, et al. 1981; Clary and Wolstenholme 1984; Peterson, et al. 2004), but the sampling of mitogenomes from diverse taxonomic groups has revealed extensive variation in complements of mt-tRNAs with examples of extreme mt-tRNA loss. Trypanosomatids such as *Leishmania tarentolae* and *Trypanosoma brucei* (Simpson, et al. 1989; Hancock and Hajduk 1990) entirely lack tRNA genes in their respective mitogenomes, and some bilaterian animals only have a single mt-tRNA gene (Lavrov and Pett 2016). In contrast, plants have an intermediate and heterogenous set of mt-tRNA genes. The mitogenomes of flowering plants typically have 11-13 native (mitochondrial in origin) tRNA genes in addition to a variable number of intracellularly and horizontally transferred tRNAs from diverse origins including plastids, bacteria, fungi, and the mitochondria of other plants (Iams, et al. 1985; Small, et al. 1999; Kubo, et al. 2000; Rice, et al. 2013; Knie, et al. 2015; Sanchez-Puerta, et al. 2019; Warren and Sloan 2020b). Despite this mosaic of mt-tRNA genes, no plant species appears to have a sufficient set in its mitogenome to decode all amino acids (Michaud, et al. 2011).

The import of nuclear-encoded tRNAs in mitochondria has been demonstrated in a handful of plant species and is assumed to compensate for the insufficient coding capacity of mt-tRNAs in the mitogenome (Ceci, et al. 1995; Kumar, et al. 1996; Brubacher-Kauffmann, et al. 1999; Duchêne and Marechal-Drouard 2001; Glover, et al. 2001). Because most plants still retain many mt-tRNA genes, they require only a specific subset of imported tRNAs for mitochondrial function. The import of nuclear-encoded tRNAs into plant mitochondria from the cytosol has been shown to be complementary rather than redundant, meaning that tRNAs corresponding to the genes functionally lost from the mitogenome are selectively imported (Kumar, et al. 1996; Brubacher-Kauffmann, et al. 1999; Salinas-Giegé, et al. 2015). This specificity of cytosolic tRNA import even extends beyond the anticodon level within isoacceptor families, as tRNA-Gly transcripts have been shown to be differentially imported or excluded in *Solanum tuberosum* mitochondria depending on the anticodon (Brubacher-Kauffmann, et al. 1999). In addition, tRNA import has been shown to be associated with translation optimization and codon usage in the green alga *Chlamydomonas reinhardtii* (Vinogradova, et al. 2009).

Heterogeneity in mt-tRNA content appears to arise over short evolutionary timeframes in plants, raising questions about the evolutionary dynamics of functional replacement of tRNAs in light of the observed import specificity. The sequencing of mitogenomes from the angiosperm genus *Silene* revealed multiple species with greatly reduced numbers of tRNA genes (Fig. 1) (Sloan, Alverson, Chuckalovcak, et al. 2012). Many of these mitochondrial gene losses appear to have happened recently as the closely related species *S. latifolia*, *S. vulgaris*, *S. noctiflora*, and *S. conica* had nine, four, three, and two mt-tRNA genes respectively (Fig. 1). These four species represent different sections within the same subgenus and have been difficult to resolve phylogenetically (Jafari, et al. 2020), making their precise history of shared and independent tRNA-gene losses unclear.

**Fig. 1.**
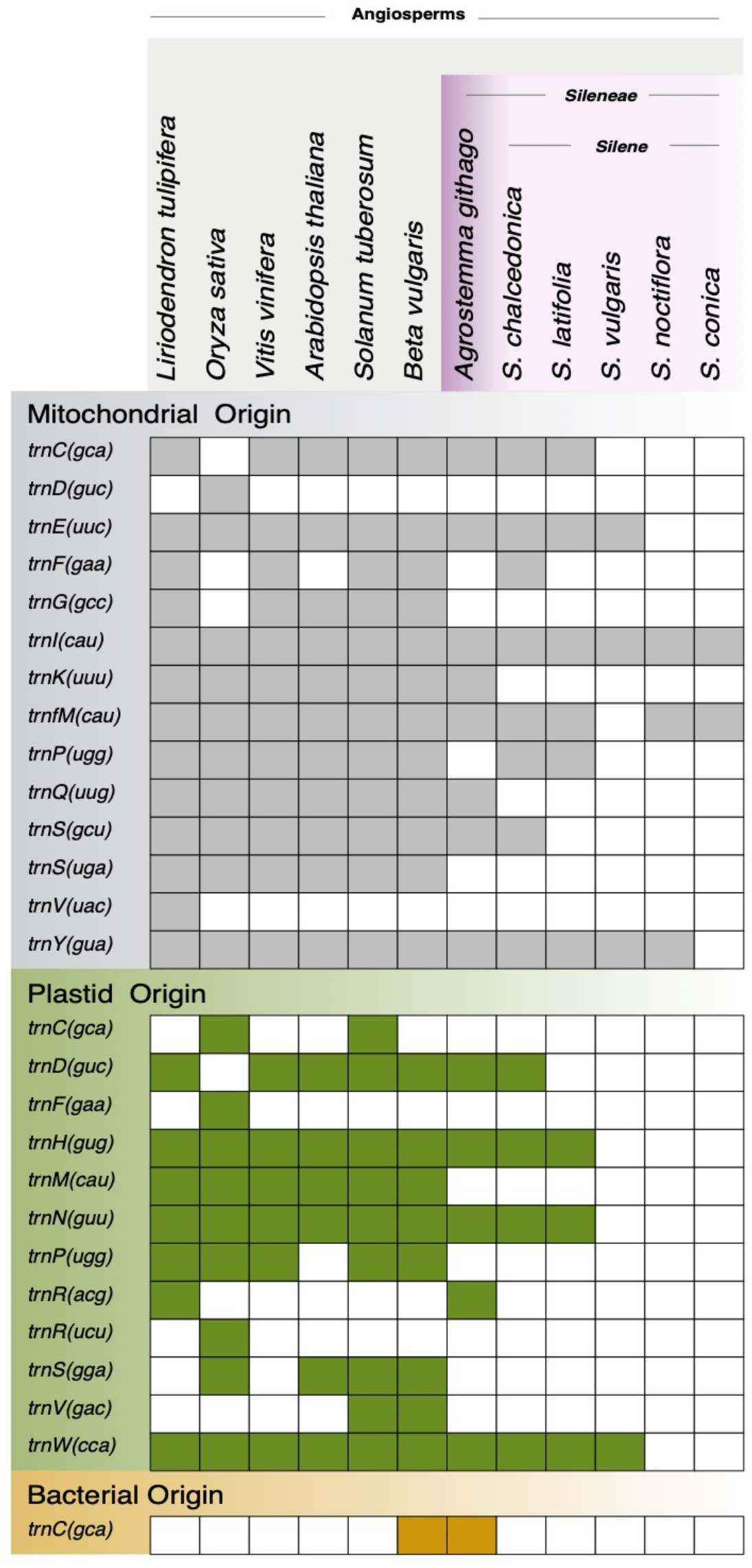
Summary of tRNA gene content in a sample of angiosperm mitogenomes with an emphasis on the family Caryophyllaceae. Filled squares indicate the presence of an intact gene sequence, with genes named based on a single-letter amino-acid abbreviation and with anticodon indicated parenthetically. Formyl-methionine is abbreviated fM. Different colors represent the ancestral origin of the tRNA gene. Disrupted or incomplete sequences inferred to be pseudogenes are not included.

This rapid and ongoing loss of mt-tRNAs in *Silene* presents a unique opportunity to study the evolutionary mechanisms involved in the functional replacement of mt-tRNAs with nuclear-encoded counterparts. However, characterizing the mt-tRNA pools (including both imported and mitochondrially encoded tRNAs) presents numerous challenges—the first being the intrinsic difficulty in sequencing tRNAs. The vast majority of RNA-seq methods require the reverse transcription of the molecules before sequencing, but the extensive base modifications in tRNAs can stall or terminate reverse transcriptase enzymes (Wilusz 2015). Additionally, the tightly base paired 5’ and 3’ termini of tRNAs can prevent the ligation of sequencing adapters (Shigematsu, et al. 2017). To date, the sequencing of purified mt-tRNA pools has only been done in humans (Mercer, et al. 2011). A second challenge in mt-tRNA genetics arises from the degenerate nature of mt-tRNAs. Although mt-tRNAs in plants generally have canonical shape (i.e. a cloverleaf-shape with a D-arm, a T-arm, an acceptor stem, an anticodon stem, and associated loops (Raina and Ibba 2014)), mt-tRNAs from other lineages (particularly metazoans) can have such aberrant structure that determining the number and location of mt-tRNAs is unreliable using only genomic data and predictive software (Boore, et al. 2005; Juhling, et al. 2009; Warren and Sloan 2020a). The existence of degenerate mt-tRNAs in some lineages raises the possibility that putatively lost tRNA genes in *Silene* are still present in the mitogenome but simply undetected. A third challenge is that the majority of tRNA import research in plants has relied on northern blot analysis (Kumar, et al. 1996; Delage, et al. 2003). This approach requires previously predicted tRNA sequences for hybridization and can lack clear resolution of probed sequences because of cross-hybridization to undesired targets. Therefore, multiple questions remain about the composition of tRNA populations in the plant mitochondrial matrix.

Here, we trace the evolutionary history of a plant lineage with extreme mt-tRNA loss using recently developed tRNA-seq methods and mitogenome sequencing. We provide the first global sequencing of tRNA pools from isolated plant mitochondria, characterizing the expression of mt-tRNA genes as well the population of imported cytosolic sequences. Using these analyses, we address the following questions: First, have *Silene* mitogenomes really experienced an extreme reduction in mt-tRNA content or do they contain “hidden” genes that are unrecognizable with standard annotation methods? If mt-tRNAs have been lost and functionally replaced by the import of cytosolic tRNAs, are certain nuclear-encoded tRNAs dedicated to mitochondrial translation? And finally, do *Silene* species preserve the import specificity that has been demonstrated in other plant species, maintaining the avoidance of functionally redundant tRNA import?

This study supports the conclusion from previous mitogenome annotations that *Silene* species have experienced extensive mt-tRNA gene loss and reveals widespread shifts in the abundance of certain nuclear-encoded tRNAs in isolated mitochondria across *Sileneae* species – suggesting that import of tRNAs from the cytosol compensates for mt-tRNA gene losses. Additionally, the functional replacement of mt-tRNAs with nuclear-encoded counterparts does not appear to involve the expression and import of tRNA genes dedicated for mitochondrial function. Finally, we show multiple instances where tRNAs with anticodons still encoded in the mitogenome appear to be imported in *Sileneae*, indicating a loss of the tRNA import specificity previously seen in other plant mitochondria.

## Results

### Tracing the evolution of mt-tRNA gene loss in *Sileneae* by sequencing the mitochondrial genomes of *Agrostemma githago* and *Silene chalcedonica*

All previously sequenced *Silene* mitogenomes have been found to have extensive mt-tRNA gene loss (Sloan, Alverson, Chuckalovcak, et al. 2012); however, all of the species that have been sequenced are from a single subgenus, *Behenantha*. In order to better reconstruct the history and timing of mt-tRNA gene losses, we expanded mitogenome sampling by sequencing a representative of the subgenus *Lychnis* (*Silene chalcedonica*) (Jafari, et al. 2020), as well as *Agrostemma githago*, which is also a member of the tribe *Sileneae* (Fig. 1). Prior to this work, the closest sequenced relative to the genus *Silene* was *Beta vulgaris* (sugar beet), representing an estimated 70 Myr of divergence from *Silene* (Magallon, et al. 2015). The mitogenome of *B. vulgaris* has a more typical set of 19 mt-tRNA genes (Kubo, et al. 2000) and lacks the history of gene loss seen in *Silene*.

The assembly of the *A. githago* mitogenome produced a 262,903 bp master circle with a GC content of 44.7%. The assembly included four copies of a 2,427 bp “core” repeat region, with additional repeated sequence flanking this core region but only present in a subset of the four copies. This structure of large, identical repeat regions is similar to the mitogenome of *S. latifolia*, which has six copies of a (different) large repeat (Sloan, et al. 2010). These repeats likely undergo intra- and intermolecular recombination, resulting in multiple genome configurations not depicted by the master circle represented here (Supp. Fig. 1).

The protein-coding gene inventory of *A. githago* is similar to that of members of the genus *Silene* (Supp. Fig. 1). Interestingly, the intron-containing gene *nad7* appears to be *trans*-spliced in *A. githago*. While the first four exons are found in a standard *cis* configuration, the fifth exon is located elsewhere in the genome (Supp. Fig. 1). Other NADH-ubiquinone oxidoreductase genes (specifically *nad1*, *nad2*, and *nad5*) typically contain *trans*-splicing introns (Chapdelaine and Bonen 1991; Knoop, et al. 1991; Wissinger, et al. 1991; Binder, et al. 1992), but this is not the case for *nad7* in most angiosperm mitogenomes. Such transitions from *cis*- to *trans*-splicing are relatively common in plant mitochondria (Qiu and Palmer 2004), and it was recently shown that the Pinaceae has independently evolved *trans*-splicing of this same *nad7* intron (Guo, et al. 2020).

The complement of 14 mt-tRNA genes in *A. githago* also falls below the typical number encoded in angiosperm mitogenomes, but it is greater than the previously documented mt-tRNA gene content in *Silene* (Fig. 1), putting *A. githago* at an intermediate state of mt-tRNA gene loss. There is, however, evidence of independent mt-tRNA gene loss in *A. githago* as both tRNA-Phe and tRNA-Pro have been lost in *A. githago* but they are present in at least one *Silene* species (Fig. 1). Two distinct copies of tRNA-Cys are present in *A. githago*: the native mitochondrial copy and the bacterial, horizontally transferred gene that was first reported in the mitogenome of *B. vulgaris* (Kubo, et al. 2000; Kitazaki, et al. 2011) (Fig. 1, Supp. Table 1). The native tRNA-Gly and tRNA-SerTGA, as well as the plastid-derived tRNA-Met and tRNA-SerGGA, appear to have been lost before the divergence of *Agrostemma* and *Silene* as they are absent from *A. githago* and all currently sequenced *Silene* species (Fig. 1). As *A. githago* does have reduced mt-tRNA gene content compared to most angiosperms, the process of tRNA gene loss likely initiated prior to divergence of *Sileneae,* but it is has proceeded to much greater extents in the sampled species from *Silene* subgenus *Behenantha*.

The mitogenome of *S. chalcedonica* was found to be considerably larger than that of *A. githago* with an estimated size of approximately 880 kb and a far more complex repeat structure. Because large, highly repetitive genomes are difficult to “close”, we did not try to produce a master circle assembly for the species. However, sequencing resulted in high coverage (>50× on average) of assembled mitochondrial contigs, suggesting that we likely captured all sequence content for identification of mt-tRNA genes. *Silene chalcedonica* was found to have 12 predicted mt-tRNA genes (two fewer than in *A. githago*). Multiple tRNA gene losses are shared by other *Silene* species (Fig. 1). For example, tRNA-Gln was likely lost before the divergence of *S. chalcedonica* and the other *Silene* species, as none of the sequenced members of the genus have intact copy of the gene. However, mt-tRNA-SerGCU and mt-tRNA-Phe are still present in *S. chalcedonica*, whereas they have been lost in the other sequenced *Silene* species (Fig. 1).

### Confirming the expression of mitochondrially encoded tRNA genes in *Sileneae* species

To validate the *in silico* annotations of mitochondrially encoded tRNA genes and characterize global mt-tRNA populations, mitochondria were isolated from leaf tissue of five *Sileneae* species (*A. githago, S. conica, S. latifolia, S. noctiflora, and S. vulgaris*) using differential centrifugation and discontinuous Percoll gradients. Paired, total-cellular RNA (whole leaf tissue) samples were processed in parallel with each mitochondrial sample. tRNAs were then sequenced with recently developed tRNA-seq protocols including treatment with the dealkylating enzyme AlkB to remove base modifications that inhibit reverse transcription (Cozen, et al. 2015; Zheng, et al. 2015) and the use of adapters with complementarity to the CCA-tail found on all mature tRNAs (Shigematsu, et al. 2017). Sequences were mapped to annotated mt-tRNA genes, as well as the entire mitogenome of each respective species to search for previously undetected tRNAs. Reads were also mapped to tRNA genes predicted from nuclear assemblies of the five species by tRNAscan-SE (Chan and Lowe 2019).

As described in detail below, the purified mitochondrial samples showed strong enrichment for mitochondrially encoded tRNAs (Fig. 2), confirming the efficacy of the mitochondrial isolations. All mt-tRNAs previously predicted to be functional in *S. conica, S. latifolia, S. noctiflora and S. vulgaris* were found to be expressed (Supp. Table 2). Two *A. githago* mt-tRNAs (tRNA-Asn, and tRNA-Arg) with identical sequence identity to plastid copies were depleted in mitochondrial isolates (Supp. Table 3), suggesting that these tRNAs may not be functional in the mitochondrial matrix. Both tRNAs have 100% sequence identity to plastid tRNAs and are present within a larger plastid-derived sequence block (i.e., a “mtpt”), indicating that these sequences are recent insertions of plastid DNA. The tRNA-Asn gene is likely different from the ancestral plastid-derived tRNA-Asn copy found in most angiosperm mitogenomes (Fig. 1), which can be differentiated from plastid copies by mitochondrial-specific sequence variants (Richardson, et al. 2013). The lack of detection of both genes suggests that that reduction in mt-tRNA gene content in *A. githago* may be more extreme than apparent from genomic data alone. Differential expression (enrichment) analysis for all mitochondrial genes can be found for each species in Supp. Tables 3-8.

**Fig. 2.**
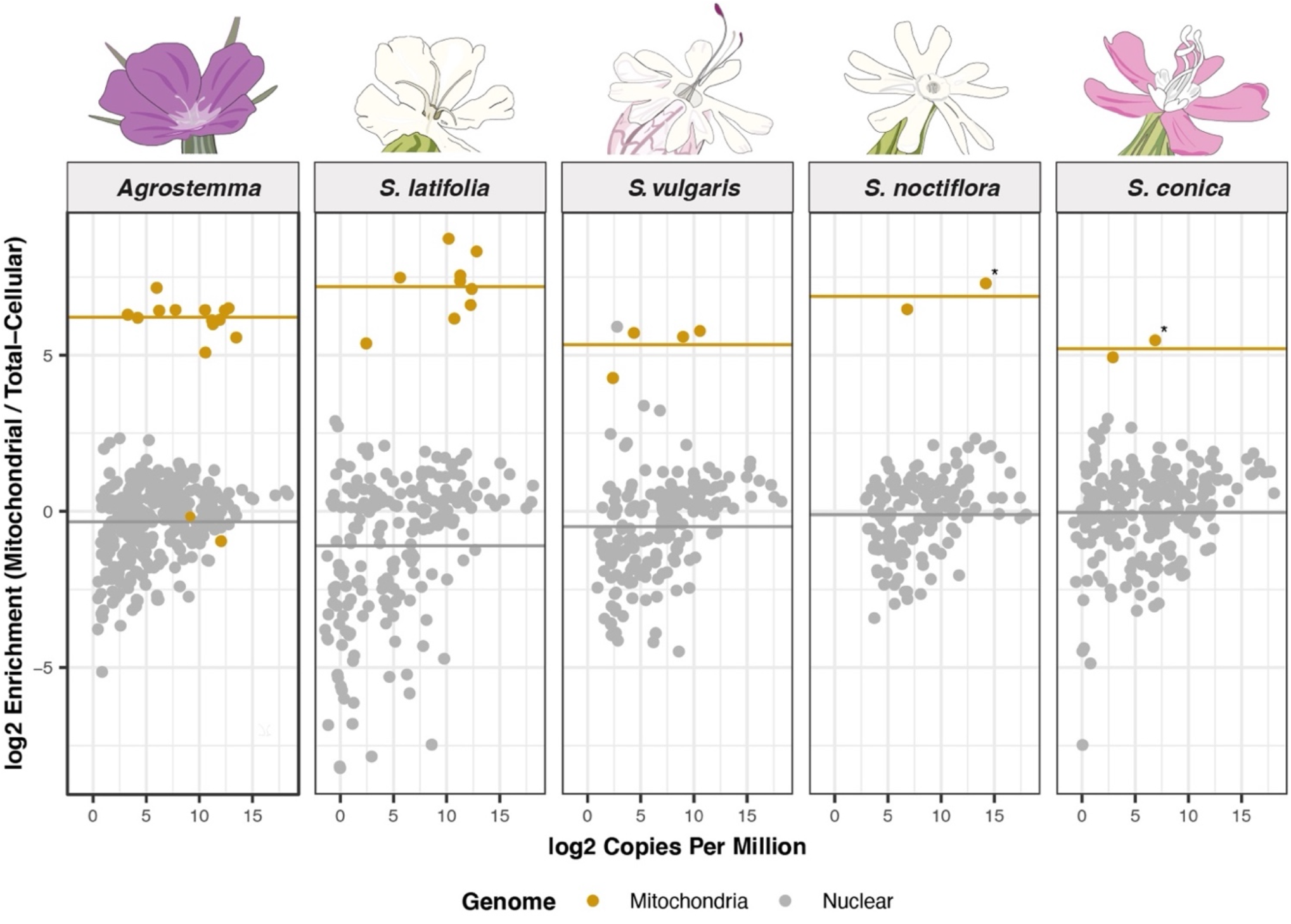
Enrichment of mitochondrially encoded and nuclear-encoded tRNAs in mitochondrial isolates relative to total-cellular samples. Each dot represents a unique tRNA sequence. The heavy gray line represents the average enrichment of nuclear-encoded genes. The genomic origin of the tRNA is indicated by color, with gray being nuclear-encoded and gold being mitochondrial. The two mt-tRNAs around the 0-line for *Agrostemma* are tRNA-Asp and tRNA-Arg, which are identical in sequence to the plastid-encoded tRNA counterparts and were not included in the calculation of average enrichment of mitochondrially-encoded tRNAs for the species. Expression of the multiple (but non-identical) copies of formyl-methionine (fMet) tRNAs present in in the mitogenomes of *S. noctiflora* and *S. conica* are marked with an * and were summed for expression analysis. Stem-loops are not shown in this figure.

### The post-transcriptional addition of a CCA tail is widespread in *Sileneae* mitochondria and occurs on multiple non-tRNA transcripts

Although the vast majority of reads from the mitogenome mapped to an annotated tRNA gene, a small percentage mapped to other positions (Table1, Supp. Table 9). These reads frequently had CCA nucleotides at a terminus and occurred at the boundaries of rRNA or protein coding genes (Supp. Table 9). For some of these sequences, the CCA nucleotides appear to be post-transcriptionally added as the sequence is not genomically encoded. Many of these reads formed stem-loop structures that act as t-element processing signals previously described in other angiosperms (Forner, et al. 2007; Varre, et al. 2019). T-elements are tRNA-like sequences bordering other genes that act as signals for endonucleolytic cleavage by RNase Z and/or RNase P for mRNA or rRNA transcript maturation (Forner, et al. 2007). A conserved t-element structure internal to the annotated *nad6* open reading frame was expressed and post transcriptionally modified with a CCA-tail in *A. githago*, *S. latifolia*, and *S. vulgaris* (Supp. Table 9, Supp. Fig. 2).

**Table 1.** Percentage of reads with a CCA-tail that mapped to a mitochondrial location other than a tRNA gene. Only sequences that had more than two reads (in all six libraries combined) were used for global mitogenome mapping and this analysis. Only two replicates of *S. latifolia* mitochondrial isolations, *S. latifolia* total-cellular samples, and *S. vulgaris* mitochondrial isolations were performed; see Methods for details.

| Species | Mitochondrial isolate 1 | Mitochondrial isolate 2 | Mitochondrial isolate 3 | Total cellular 1 | Total cellular 2 | Total cellular 3 |
| --- | --- | --- | --- | --- | --- | --- |
| *Agrostemma* | 0.2316% | 0.3290% | 0.2057% | 0.0085% | 0.0072% | 0.0086% |
| *S. latifolia* | 0.0700% | 0.0438% | NA | 0.0009% | 0.0003% | NA |
| *S. vulgaris* | 4.4819% | NA | 3.6261% | 0.1084% | 0.1553% | 0.3070% |
| *S. noctiflora* | 0.0243% | 0.0314% | 0.0234% | 0.0012% | 0.0018% | 0.0059% |
| *S. conica* | 0.0005% | 0.0008% | 0.0014% | 0.0000% | 0.0000% | 0.0000% |

Although most of these stem-loops lacked canonical tRNA structure, they did have stem structure at the 5′- and 3′-termini (Supp. Fig. 2). The CCA-tailing of stem-loop structures has been reported in numerous other mitochondrial systems including rat (Yu, et al. 2008), human (Pawar, et al. 2019), mites (Warren and Sloan 2020a), and other plants (Williams, et al. 2000). Interestingly, the species with the fewest mt-tRNAs (*S. conica* and *S. noctiflora*) also had the fewest of these detected stem-loops (Table 1, Supp. Table 9). Only 0.0005-0.0314% of the CCA-containing reads in these two species mapped to a non-tRNA position in the mitogenome, and these reads almost exclusively originated from the boundaries of rRNA genes (Supp. Table 9). In comparison, the percentage of reads mapping to non-tRNA position in the mitogenomes in the other three species ranged from 0.0438% up to 4.4819%. The relatively abundant reads mapping to a stem-loop structure in *S. vulgaris* derived mainly from a region of the mitogenome with two identical step-loop sequences that were not immediately up- or downstream of an annotated gene (over 6 kb away from *cob* and over 4.5 kb away from the fourth exon of *nad5*). *Silene vulgaris* also had a relatively highly abundant stem-loop structure originating from a tRNA-fMet gene (over 51,000 reads combined in all six libraries). This sequence is annotated as a pseudogene as it lacks multiple features of a tRNA, and it is predicted to form an arm-less, degenerate stem-loop structure with multiple bulges (Supp. Fig. 2). It is only 58 bp away from *atp6*, so it is likely that it now serves as a t-element maturation signal.

One stem-loop structure mapped to the mitogenome of *A. githago* did have a cloverleaf-like structure similar to a canonical tRNA shape but did not correspond to an annotated tRNA gene (Fig. 3, Supp. Table 2a). Predictive folding under a maximum free energy model puts the anticodon as tRNA-LysUUU (Fig. 3). Unlike t-elements, this stem-loop is not closely associated with any annotated gene, as it is 2,414 bp upstream of the nearest gene (the fourth exon of *nad1*, Supp. Table 9). Homologous sequences are present in the mitogenome of *Beta vulgaris* (GenBank: BA000024) and *Chenopodium quinoa* (GenBank: NC_041093), as well as copies located within a region of mitochondrial-like sequence in the nuclear genome assembly of *Linum usitatissimum* (GenBank: CP027627). However, this sequence is not present in the mitogenomes of any of the sequenced *Silene* species. Although the evolution of novel tRNAs through the duplication of existing tRNA genes has been shown phylogenetically and experimentally (Ayan, et al. 2020), this sequence in *Agrostemma* does not have identifiable homology to any known tRNA and does not appear to form a tRNA-like structure in *B. vulgaris* or *C. quinoa*. Whether this is just a non-functional stem-loop that happened to converge on a predicted cloverleaf structure or a functional decoding molecule representing the *de novo* birth of a tRNA gene is not clear.

**Fig. 3.**
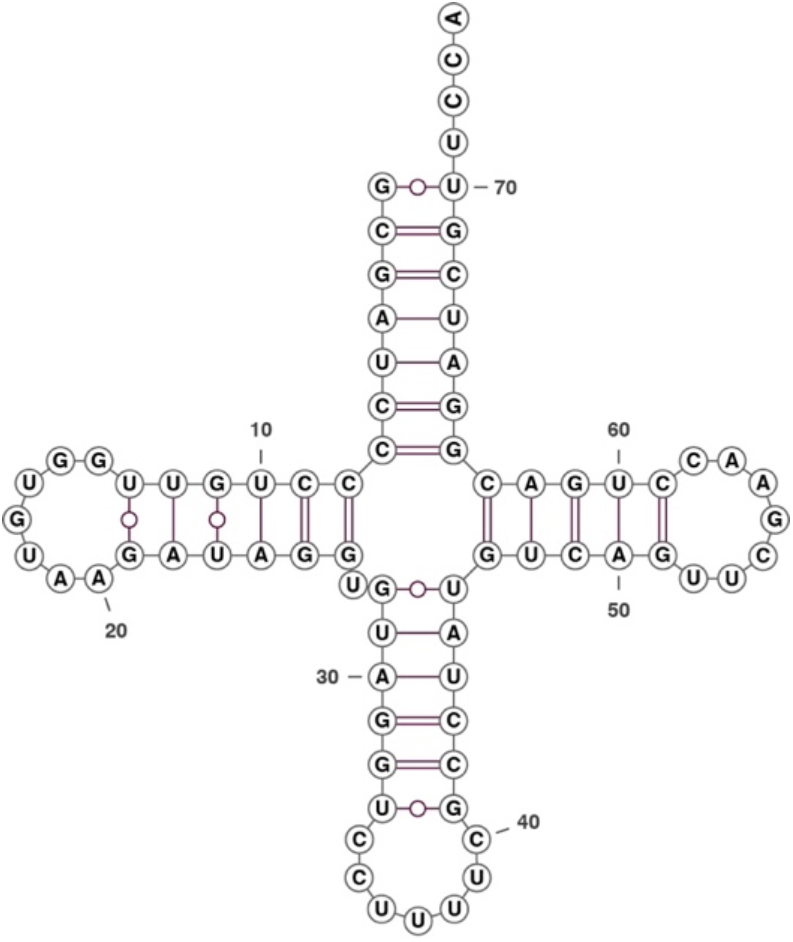
Predicted cloverleaf folding transcript expressed and post-transcriptionally modified with a CCA-tail in *A. githago*. Despite the predicted tRNA-like secondary structure, this sequence is not homologous to any previously annotated mt-tRNA genes. Folding represents the maximum free energy folding model using the program the program RNAfold (Lorenz, et al. 2011), and structure diagrams were created using the program VARNA ver. 3.9 (Darty, et al. 2009).

For multiple reasons, it is unlikely that the vast majority of the “stem-loops” detected in this dataset represent functional tRNAs. First, with the exception of the tRNA-like Lys structure in *A. githago*, these stem-loops have structural deformities that would be generally inconsistent with tRNA function, including multiple mismatches, a lack of a cloverleaf or L-shape, and incorrectly sized anticodon loops. Even in extremely aberrant metazoan mt-tRNAs that lack arms, an L-shape tertiary structure is still expected (Watanabe, et al. 2014; Juhling, et al. 2018). Mismatched bases in stems of plant mt-tRNAs have been predicted from genomic data but found to be corrected post-transcriptionally (Marechal-Drouard, et al. 1993; Binder, et al. 1994). The sequenced stem-loops found in this study maintained these base mismatches. Secondly, many of these structures originate from known t-element regions or are directly up or downstream of other genes. T-elements are broadly present in angiosperms (including those with much larger mt-tRNA gene sets) and are known substrates for other tRNA-processing machinery (i.e., RNase P/Z) (Forner, et al. 2007). Thus, it is likely that these sequences are just processed with a CCA-tail as a byproduct. Lastly, these stem-loop sequences were of low abundance and almost absent in the species with the fewest mt-tRNA genes. Therefore, it does not appear that the apparent mt-tRNA gene loss seen in *Sileneae* species can be explained by a large class of previously undetected tRNA genes in the mitogenome.

### Characterization of imported tRNA pools in *Sileneae* mitochondria reveals extensive sharing of the same nuclear-encoded tRNAs between the cytosol and the mitochondria

The average enrichment of mitochondrial-encoded tRNAs in purified *Sileneae* mitochondrial fractions was anywhere from 37- to 146-fold relative to total-cellular samples (Supp. Tables 3-8) depending on the species. Almost no nuclear-encoded tRNAs in any of the five *Sileneae* species reached the enrichment level of the mitochondrially-encoded tRNAs (Fig. 2), suggesting that very few, if any, nuclear-encoded tRNAs exclusively function in the mitochondria. Therefore, nuclear-encoded tRNA that are imported into the mitochondria must also be present in substantial abundances in the cytosol or elsewhere in the cell. The most notable exception was a low-abundance nuclear-encoded tRNA in *S. vulgaris* with homology to tRNA-AlaCGC. This tRNA was part of a family of related genes (Supp. Table 1, Reference ID#s 964, 1005-1014) with multiple sequence variants (Fig. 4) and reached comparable enrichment levels to those observed for mitochondrial genes (Fig. 2). The anticodon of these tRNAs is uncertain, as an insertion in the anticodon loop may change the anticodon sequence from CGC to ACG (tRNA-Arg) or possibly GCA (tRNA-Cys), depending on how the tRNA folds *in vivo* (Fig. 4, Supp. Fig. 4). Multiple tRNAs originating from these genes were found to be enriched in mitochondrial isolates, but detection was low and only one (nuclear ID: 1009, Supp. Table 1) survived the minimum read count threshold for differential expression analysis (Fig. 2, Supp. Table 8). Both tRNA-Arg and tRNA-Ala are expected to be imported into all angiosperm mitochondria (Michaud, et al. 2011), and tRNA-Cys is expected to be imported into *S. vulgaris* because of recent loss of a mt-tRNA-Cys gene, but why enrichment of these noncanonical tRNAs would be so high in mitochondrial isolates is unclear. It is possible that this is not a case of exclusive import but instead a failure of quality control, as import may have occurred prior to degradation, and the cytosolic surveillance mechanisms that degrade tRNAs without proper structures (Dewe, et al. 2012) may not be active in the mitochondrial matrix. Other explanations for this enrichment are possible, but regardless these genes were the exception, as there was a clear distinction in the enrichment of mt-tRNAs versus the nuclear-encoded tRNA populations (Fig. 2).

**Fig. 4.**
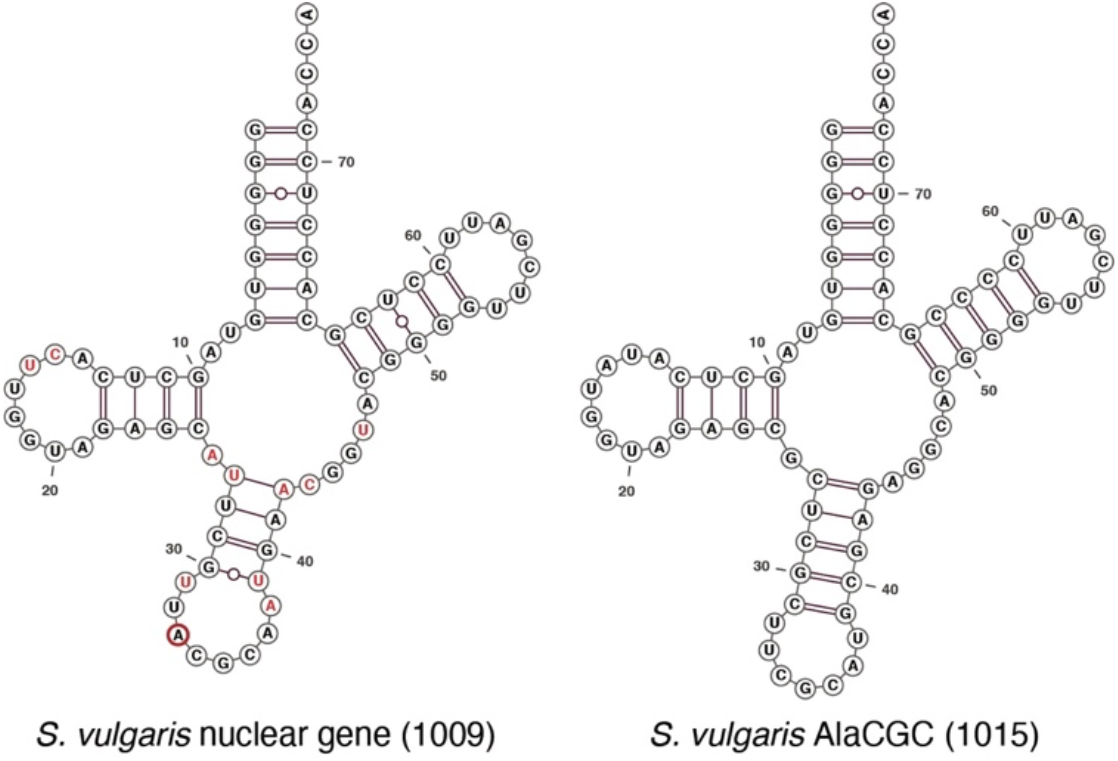
Sequence and predictive folding of a highly enriched nuclear-encoded gene (#1009) in *S. vulgaris* mitochondrial isolates and the closest *S. vulgaris* reference with a tRNAscan-determined anticodon. The unique reference ID is in parentheses. Red letters indicate base pair differences between the two tRNAs, and the thick, red circle indicates the insertion in the anticodon mentioned in the main text. Nuclear gene 1009 had multiple more closely related sequences but without definitive anticodons see Supp. Table 1 nuclear reference ID#s 964, 1005-1014.

### Confirming the utility of tRNA-seq methods to detect the import of nuclear-encoded tRNAs by sequencing mitochondrial tRNA pools in *Solanum tuberosum*

Because *S. tuberosum* (potato) has had the most extensive prior characterization of mitochondrial tRNA import using northern blot analysis (Marechal-Drouard, et al. 1990; Kumar, et al. 1996; Brubacher-Kauffmann, et al. 1999; Delage, et al. 2003; Pujol, et al. 2008), we sequenced the mitochondrial tRNA pools in *S. tuberosum* in order to compare the effectiveness of detecting tRNA import using the tRNA-seq methods employed in this study. Mitochondria were isolated from young *S. tuberosum* leaves with the same protocol and Percoll gradients used for *Sileneae* species.

Enrichment of *S. tuberosum* mitochondria in the isolated fractions was apparent from high (21-fold-change) relative abundance of mt-tRNA and mitochondrial stem-loop sequences compared to the total-cellular samples (Supp. Fig. 3, Supp. Table 6). Likewise, plastid tRNAs were greatly depleted relative to mitochondrial tRNAs, indicating that the preparation of the mitochondrial isolates was also effective (although not perfect) at removing plastid tRNAs (Table 2). As there is no known import of plastid tRNAs into mitochondria, any detection of plastid tRNAs in mitochondrial fractions likely represents contaminating plastids in the mitochondrial gradient purification.

**Table 2.** Depletion of tRNAs from the nuclear and plastid genomes in mitochondrial isolates from six angiosperm species relative to mitochondrially encoded tRNAs. Nuclear tRNA depletion was calculated as the difference between the average log_2_ fold change of all mt-tRNA genes (including mitochondrial stem-loops) and the average log_2_ fold change of nuclear-encoded tRNAs. Average plastid tRNA depletion was calculated similarly as the difference between the average mt-tRNA log_2_ fold change and the average log_2_ fold change of plastid tRNAs. ^*^Note that the depletion values for nuclear-encoded tRNAs do not account for the fact that some of these tRNAs are presumably imported into the mitochondria. Thus, the values reported here likely underestimate the true depletion levels for non-imported, nuclear-encoded tRNAs.

|  | **Average plastid tRNA depletion** | **Average nuclear tRNA depletion\*** |
| --- | --- | --- |
| **Species** | Relative log<sub>2</sub> fold difference | Relative log<sub>2</sub> fold difference |
| *Solanum* | -4.43 | -4.75 |
| *Agrostemma* | -6.76 | -6.55 |
| *S. latifolia* | -7.56 | -8.57 |
| *S. vulgaris* | -6.37 | -5.78 |
| *S. noctiflora* | -8.68 | -6.99 |
| *S. conica* | -7.45 | -4.88 |

Previous experiments testing the exclusion of cytosolic tRNAs in *S. tuberosum* (Marechal-Drouard, et al. 1990; Brubacher-Kauffmann, et al. 1999) reported a strong exclusion of tRNAs with the same anticodon as those that are still encoded in the mitogenome. In order to compare the sequence data generated here to previously performed hybridization methods (which detects the abundance of multiple tRNAs with the same anticodon), we calculated an average mitochondrial enrichment (log_2_ fold-change) weighted by expression of all tRNA sequences sharing the same anticodon (Supp. Table 10). The resulting mitochondrial enrichment values for tRNAs grouped by anticodon were highly consistent with previous northern blot results on enrichment and depletion of nuclear-encoded tRNAs in the mitochondrial fraction (Supp. Table 11). The seven tRNAs previously shown to be imported almost universally had higher enrichment values than the 17 tRNAs previously shown to be excluded – the only exception being that that tRNA-Gln(TTG) (excluded) had a slightly higher enrichment value than tRNA-Thr(TGT) (imported) (Supp. Table 11). The support for expected import and exclusion patterns for cytosolic tRNAs in *S. tuberosum* included clear differential import of tRNA-Gly(CCC) and tRNA-Gly(UCC) transcripts but exclusion of tRNA-Gly(GCC) (Supp. Table 11) (Brubacher-Kauffmann, et al. 1999).

Because only a subset of *S. tuberosum* tRNAs have been previously assayed for mitochondrial import by northern blot, we extended our analysis of the tRNA-seq data by predicting import/exclusion status for the remaining anticodons. An isodecoder family was predicted to be excluded if a tRNA for that anticodon was still encoded in the *S. tuberosum* mitogenome or imported if there was no corresponding tRNA encoded in the mitogenome. Correspondence between tRNA-seq enrichment/depletion and expected import status was highly significant (i.e., tRNAs that were predicted to be imported were significantly more enriched in the mitochondrial fraction than those expected to be excluded; Welch’s t-test, *p*-value = 3.243e-07 Fig. 5).

**Fig. 5.**
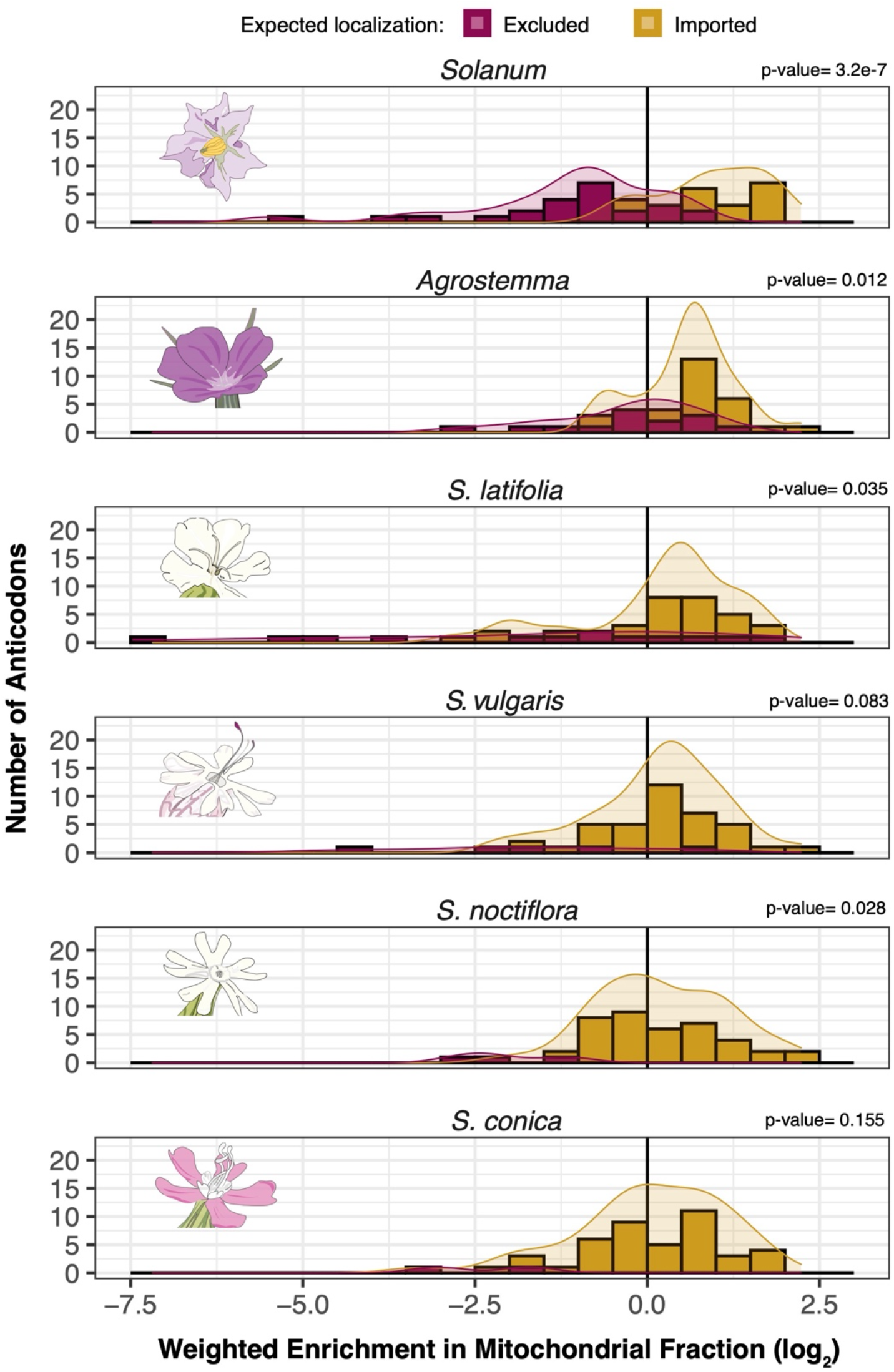
Histograms and density curves showing the distribution of average mitochondrial enrichments of nuclear-encoded tRNA anticodon families in *S. tuberosum, A. githago* and four *Silene* species. tRNAs with anticodons that are expected to be imported due to the lack of a corresponding mt-tRNA gene are shown in gold. Those that are expected to be excluded are shown in magenta. Reported *p*-values are based on *t*-tests between these two groups. Enrichment values are calculated based on expression-weighted average of all tRNAs with the same anticodon (see Methods). Positive and negative values indicate enrichment and depletion in the mitochondrial fraction, respectively.

### Extensive shifts in mitochondrial import and exclusion of tRNAs in Sileneae with possible cases of redundant tRNA import

We wanted to test whether rapid mt-tRNA gene loss in *Sileneae* precipitated shifts in cytosolic tRNA trafficking through changes in the import and exclusion of certain tRNA families. Like in *Solanum*, a signal of tRNA exclusion and import was also recovered for *Sileneae* species, with tRNA exclusion corresponding to some anticodons retained in the mitogenomes (i.e., a signal of non-redundant import of nuclear-encoded tRNAs). For all five *Sileneae* species, cytosolic tRNAs that were predicted to be excluded from mitochondria because they share an anticodon with a mitochondrially encoded tRNA showed lower relative abundance on average than those predicted to be imported (Fig. 5). This difference was not statistically significant for *S. conica* and *S. vulgaris* (Fig. 5), but that may simply reflect the limited statistical power in species where so few tRNAs are predicted to be excluded because of extensive tRNA gene loss from the mitogenome. Even in these species, the cytosolic tRNAs that corresponded to the few remaining anticodons remaining in the mitogenomes were often the most depleted tRNAs (Supp. Tables 3-8, and Supp. Table 10). For example, cytosolic tRNA-Ile(TAT) was always one of the most strongly excluded tRNA families in all species, and the functionally equivalent mt-tRNA-Ile(CAT) is the only mt-tRNA gene retained in all species tested in this analysis (Fig. 1).

We predicted that *Sileneae* species that had lost a mt-tRNA gene would exhibit evidence of increased import of the corresponding nuclear-encoded tRNA from the cytosol. Overall, of the 12 mt-tRNA genes lost in at least one *Silene* species since divergence from *Agrostemma*, seven corresponding cytosolic tRNAs (tRNA-Asp(GTC), tRNA-Cys(GCA), tRNA-Gln(TTG), tRNA-Glu(TTC), tRNA-His(GTG), tRNA-Trp(CCA) and tRNA-Tyr(GTA)) showed increased representation in the mitochondrial fraction in all species that had lost the mt-tRNA counterpart compared to both *Solanum* and *Agrostemma* (Figs. 6-7, Supp. Fig 4). Conversely, there were no cases showing the opposite pattern, where all *Silene* species with a mt-tRNA gene loss had lower enrichment than both *Solanum* and *Agrostemma* for the corresponding cytosolic tRNA. In addition, there are two mt-tRNA genes that were lost in the *Sileneae* lineage prior to the divergence of *Agrostemma* and *Silene*: tRNA-Gly(GCC) and tRNA-Met(CAT) (Fig. 1). In both of these cases, the corresponding cytosolic isodecoder families showed higher mitochondrial enrichment in all five *Sileneae* species than *Solanum* (Supp. Fig. 5). In contrast to these tRNAs with a recent history of loss of the corresponding mt-tRNA genes in *Silene*, the nuclear-encoded tRNAs that are expected to be universally imported in angiosperms because of more ancient losses from the mitogenome did not show consistent trends towards mitochondrial enrichment in *Silene* relative to *Agrostemma* and *Solanum* (Supp. Fig. 5)

**Fig. 6.**
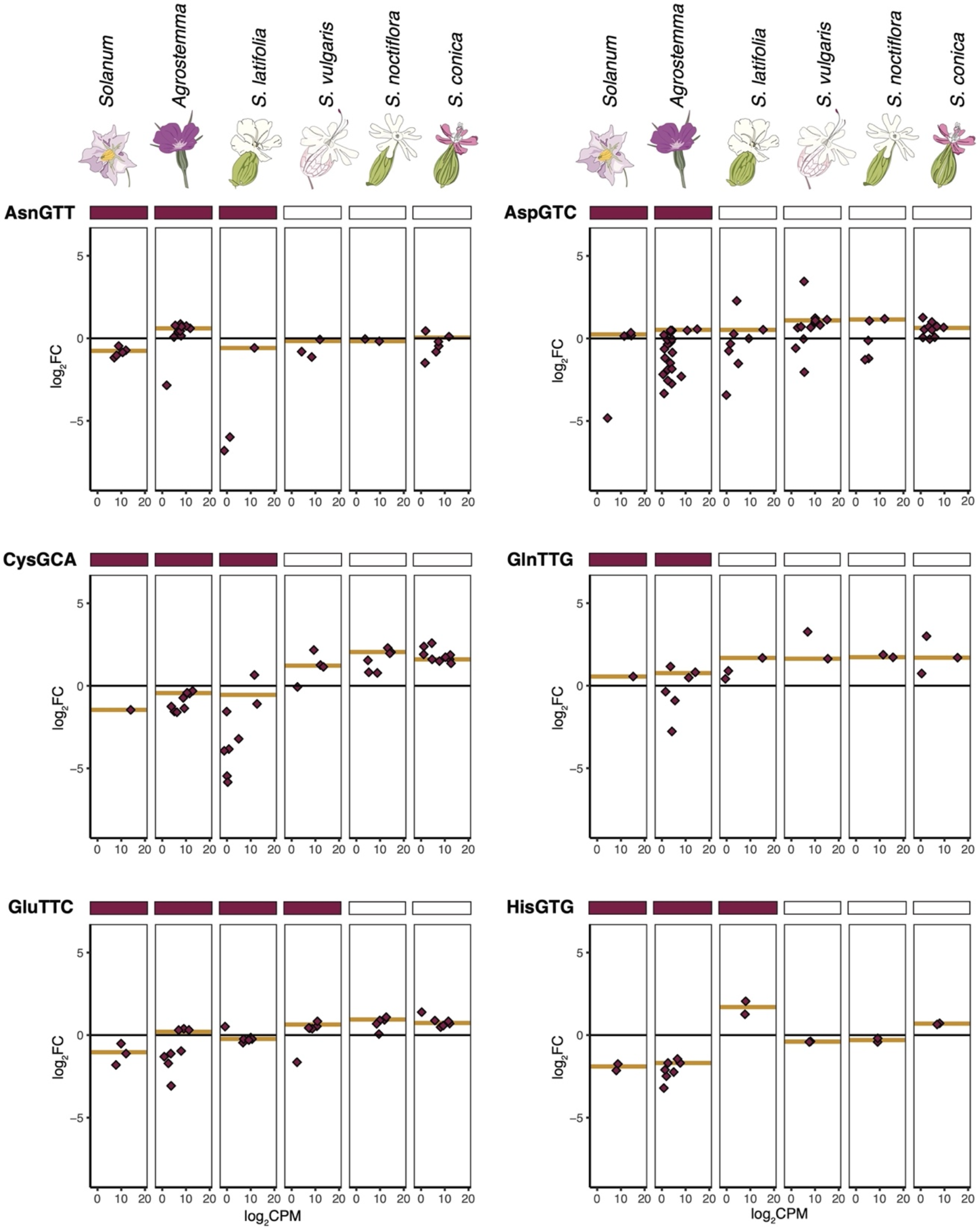
Enrichment and expression of individual cytosolic tRNA genes corresponding to mitochondrial genes with a history of loss in *Silene* since divergence from *Agrostemma* (first half, see Fig. 7 for second half). The y-axis is enrichment in log_2_ fold change in mitochondrial isolates versus total-cellular samples. The x-axis is expression level in counts per million on a log_2_ scale. Points represent individual reference sequences, and gold lines represent the average enrichment of all nuclear-encoded tRNAs with that anticodon weighted by expression (see Methods; enrichment data for all anticodons can be found in Supp. Table 10). A filled rectangle above a panel indicates that a tRNA gene with that anticodon is encoded in the mitogenome of that species.

**Fig. 7.**
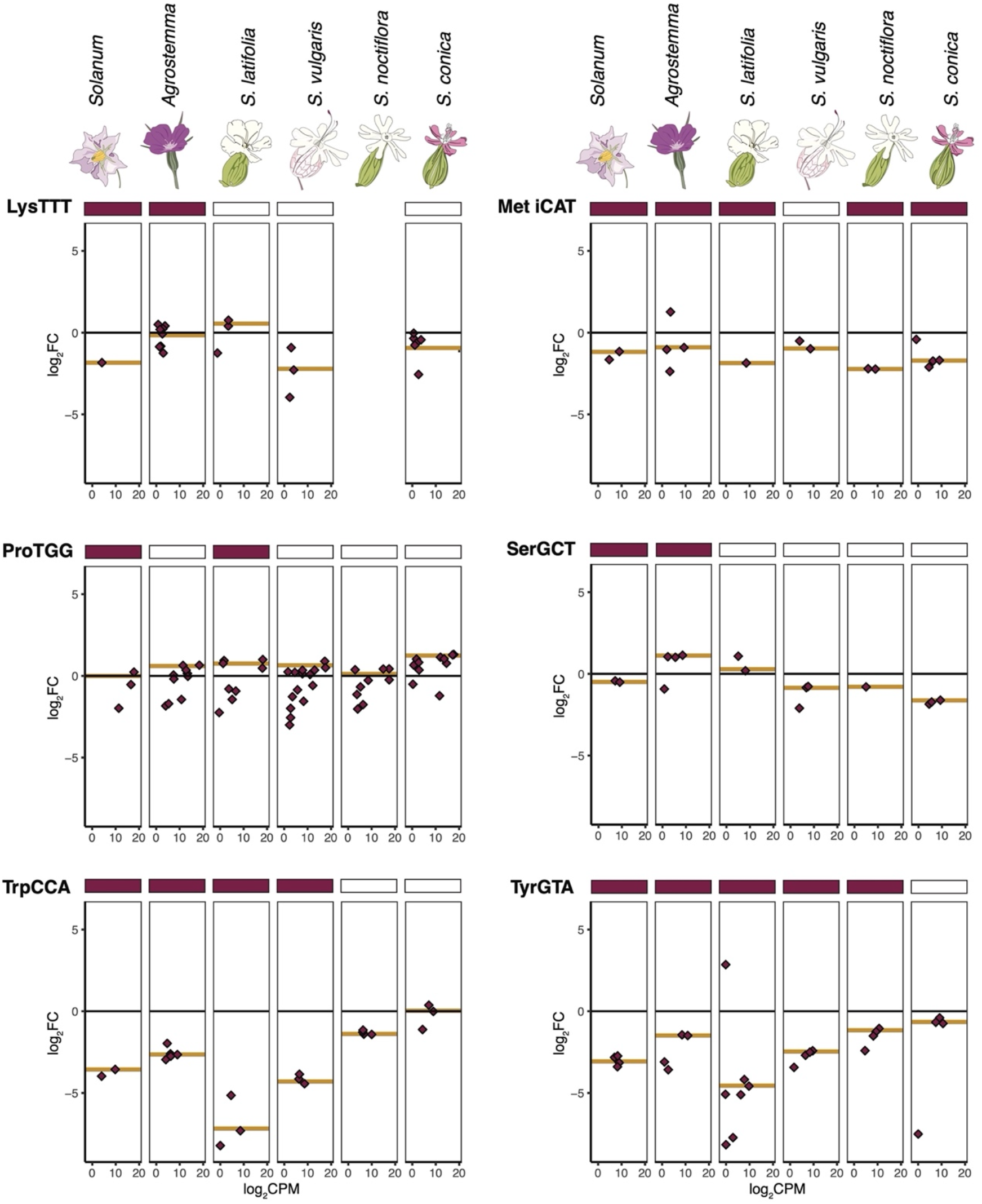
Enrichment and expression of individual cytosolic tRNA genes corresponding to mitochondrial genes with a history of loss in *Silene* since divergence from *Agrostemma* (second half, see Fig. 6 for first half). The y-axis is enrichment in log_2_ fold change in mitochondrial isolates versus total-cellular samples. The x-axis is expression level in counts per million on a log_2_ scale. Points represent individual reference sequences, and gold lines represent the average enrichment of all nuclear-encoded tRNAs with that anticodon weighted by expression (see Methods; enrichment data for all anticodons can be found in Supp. Table 10). A filled rectangle above a panel indicates that a tRNA gene with that anticodon is encoded in the mitogenome of that species. No panel is reported for *S. noctiflora* tRNA-Lys because no tRNA-Lys gene was detected as sufficient expression levels of analysis of differential expression in that species.

In some cases of mt-tRNA gene loss, we did not see expected changes in import. For example, the cytosolic initiator tRNA-Met did not exhibit positive enrichment in any species (Supp. Fig. 4), including *S. vulgaris* which lacks and intact copy of the functionally equivalent tRNA-fMet from its mitogenome – possibly suggesting that initiator tRNA-Met may not be used for mitochondrial translation initiation even in systems that lose tRNA-fMet. Multiple tRNA-Ser anticodons failed to show a consistent pattern of increased enrichment despite multiple losses of tRNA-Ser genes in *Sileneae* mitogenomes but did exhibit substantial heterogeneity in enrichment across the different species (Supp. Fig. 5).

Interestingly, there were multiple cases of increased mitochondrial enrichment of a cytosolic tRNA in a *Sileneae* species that still retained the corresponding mt-tRNA gene. Such cases suggest that there may be sustained periods of time where mitochondrially encoded tRNAs and imported cytosolic tRNAs with the same anticodon exist redundantly within mitochondria. For tRNA-His(GTG) in *S. latifolia*, tRNA-Glu(TTC) in *S. vulgaris*, and tRNA-Asn(GTT) in *A. githago* (Figs. 6–7, Supp. Fig. 4 and 5), there was a positive enrichment value despite the species retaining a mitochondrial counterpart in the mitogenome.

## Discussion

The import of tRNAs across mitochondrial membranes occurs in various eukaryotic lineages (Salinas-Giegé, et al. 2015), but questions remain about the evolutionary mechanisms that facilitate the functional replacement and integration of nuclear-encoded tRNAs in mitochondrial translation.

Except for a handful of tRNAs related to tRNA-AlaCGC in *S. vulgaris*, we did not find nuclear-encoded tRNAs enriched to the same level as those encoded in the mitogenome (Fig. 2). This result suggests an absence of dedicated tRNAs for mitochondrial import in these systems with recent mitochondrial tRNA gene replacement. The widespread sharing of nuclear-encoded tRNAs between both compartments is notably different than the imported mitochondrial proteome, which has been estimated to only be 15% dual-localized between the mitochondria and another cellular location (Calvo and Mootha 2010). This lack of tRNA specialization contrasts with mitochondrial tRNA import studies performed in the green alga *Chlamydomonas* in which multiple nuclear-encoded tRNAs had varying degrees of mitochondrial specialization, and a nuclear-encoded tRNA-Lys(UUU) was found to be exclusively localized to the mitochondria (Vinogradova, et al. 2009).

One caveat to this interpretation is that, even if tRNAs are destined for exclusive function in the mitochondria, there may be a substantial fraction “en route” at any given point in time. Therefore, they would still be present in substantial quantities elsewhere in the cell, producing a muted mitochondrial enrichment signal. This caveat aside, the dual localization of tRNAs between the cytosol and mitochondria represents a large-scale homogenization of host (nuclear) and endosymbiont (mitochondrial) translational systems.

Sequencing tRNA pools from isolated mitochondria of multiple *Sileneae* species revealed substantial changes in the relative abundance of nuclear-encoded tRNAs corresponding to mt-tRNA gene loss (Figs. 6–7 and Supp. Fig. 4). These fluctuations occurred on phylogenetically recent timeframes, suggesting that changes to tRNA import specificity can occur rapidly. However, the *de novo* evolution of tRNA import did not appear to be a perfectly binary switch from excluded to imported. There were cases where a nuclear-encoded tRNA family showed partially elevated abundance in the mitochondrial fraction of a species that still retained a mitochondrially encoded counterpart (see tRNA-Glu and tRNA-Tyr in Fig. 6 and Fig. 7). Loss of exclusion signals prior to mt-tRNA gene loss may point to a model where the evolution of tRNA import is gradual process whereby some cytosolic “leakiness” is tolerated. If these imported tRNAs are functional, we hypothesize that leaky tRNA import may provide the necessary first step in tolerating mt-tRNA gene losses by providing functional redundancy. A weak signal of redundant import of a cytosolic tRNA has been previously reported in *S. tuberosum* and *Triticum tivum* (Marechal-Drouard, et al. 1990; Glover, et al. 2001), and similarly proposed to be functional alongside the mitochondrial counterpart (Glover, et al. 2001).

And finally, it is interesting to note that some of the mt-tRNAs most recalcitrant to loss in *Silene* (tRNA-Ile, tRNA-Trp, and tRNA-Tyr) corresponded to the most excluded cytosolic tRNAs in *Solanum*. It may be that certain tRNAs interact with additional/different enzymatic partners during cellular trafficking, thereby preventing indiscriminate import and resulting in strong exclusion signals. The successful delivery of these tRNAs to functionally sufficient levels would then require additional coevolutionary hurdles to overcome before functional replacement.

tRNAs are not standalone molecules in the process of translation, as entire networks of enzymes must interact with tRNAs for maturation and function. Because some tRNA-interacting enzymes are highly specific (Chimnaronk, et al. 2005; Salinas-Giegé, et al. 2015), separate sets of enzymes are normally required for the maturation and aminoacylation of cytosolic vs. organellar tRNAs (Duchêne, et al. 2009). How nuclear-encoded tRNAs are effectively “swapped” into the mitochondrial translational machinery has remained a long-standing question in plant mitochondrial genetics (Warren and Sloan 2020b). Notably, the charging of tRNAs with the correct amino acid is necessary for protein synthesis and relies on aaRSs, which use specific nucleotide identities along the tRNA for recognition and interaction with cognate tRNAs (Giegé, et al. 1998; Meiklejohn, et al. 2013). Without an effective aaRS partner, a newly imported tRNA would be nonfunctional in translation. One possible evolutionary mechanism would involve a cytosolic aaRS also gaining mitochondrial import and charging the cognate, imported tRNA present in the mitochondrial matrix. Massive tRNA loss in the mitogenomes of some non-bilaterian animals (e.g., sponges) has been associated with the loss of mitochondrial aaRSs, suggesting that imported tRNAs are being charged by partnered cytosolic aaRSs (i.e., replacement of both tRNA and aaRS) (Haen, et al. 2010; Pett and Lavrov 2015). However, this presents what has been previously termed the “chicken and egg” problem in plant mt-tRNA replacement (Small, et al. 1999). Does the aaRS or the tRNA gain import first if nether would be functional without the other? Our results from sequencing of tRNAs from the mitochondrial fractions of *Sileneae* species as well as *S. tuberosum* may provide some insight into this conundrum—cytosolic tRNAs not essential for mitochondrial translation (because of the presence and expression of a native mt-tRNA counterpart) may frequently be found in the mitochondrial matrix before the loss of mt-tRNA genes. It is yet to be determined if these imported tRNAs are functional, and if so, whether a mitochondrial aaRSs has evolved increased substrate recognition to charge both mitochondrial- and nuclear-encoded tRNAs, or if cytosolic aaRSs has also gained mitochondrial import alongside the cognate tRNA.

The homogenization of tRNA metabolism in plant mitochondria represents a much different evolutionary route than that taken by most bilaterian animals, which have retained an independent and conserved set mt-tRNAs with very limited tRNA import into mitochondria (Grosjean and Westhof 2016). Fundamental differences in the historical mutation rates of mitogenomes in animals versus plants may have created a situation whereby plant mt-tRNAs are more predisposed to the functional replacement with that of cytosolic counterparts. The rate of mitochondrial sequence evolution in land plants are among some of the slowest ever estimated (Mower, et al. 2007) and has likely been a factor in the maintenance of the largely canonical shape of plant mt-tRNAs. This conserved shape may contribute to interchangeable nature of plant mt-tRNA genes. It is possible that slow rate of evolution in plant mitogenomes has resulted in tRNA enzymatic networks that are more amenable to nuclear-encoded substrates because of shared sequence identity and structure between plant mt-tRNAs and nuclear counterparts. In contrast, the much faster evolving animal mitogenomes (Brown, et al. 1979; Lavrov and Pett 2016) may have experienced early tRNA mutations and compensatory enzymatic substitutions that make nuclear-encoded tRNAs poor substrates for the tRNA-interacting enzymes that function in the mitochondria (Watanabe, et al. 2014; Pons, et al. 2019) — thereby locking in independence of the mitochondrial protein synthesis apparatus.

Despite this generally low mutation rate, some plant lineages have experienced a recent and dramatic acceleration in mutation rates in their mitogenomes, including multiple species in *Silene* (Mower, et al. 2007; Sloan, et al. 2009; Broz, et al. 2021). This accelerated rate may have then facilitated a rapid replacement of mt-tRNA genes in some plant lineages. *Silene conica* and *S. noctiflora* both have highly accelerated rates of mitochondrial sequence evolution, as well as the fewest reported mt-tRNA genes in angiosperms, with only two and three mt-tRNA genes, respectively (Fig. 1). It has been proposed that high mutation rates increase the likelihood of disruptive mutations in tRNAs and provide a selective pressure to functionally replace mt-tRNA genes with imported nuclear-encoded counterparts – thereby ratcheting a gene replacement process (Brandvain and Wade 2009). Another plant lineage with a high rate of sequence evolution, *Viscum* (mistletoe), also has a severely reduced mt-tRNA gene content (Petersen, et al. 2015; Skippington, et al. 2015). The heterogeneity in mitogenomes across angiosperm diversity creates exciting opportunities to test these hypothesized relationships between import specificity, mutation rates, and the functional loss/replacement of mt-tRNAs.

## Methods

### Agrostemma githago *and* Silene chalcedonica *mitochondrial genome sequencing*

Mitochondrial DNA was extracted from *A. githago* and *S. chalcedonica* using methods described previously (Sloan, et al. 2010; Sloan, Alverson, Wu, et al. 2012). Tissue was collected from the same maternal families previously used for plastid genome sequencing in these species (Sloan, et al. 2014). Mitochondrial DNA from each species was used to construct 3kb paired-end libraries which were sequenced on a Roche 454 GS-FLX platform with Titanium reagents. Library construction and sequencing were performed at the University of Virginia’s Genomics Core Facility. Reads were assembled with Roche’s GS de novo Assembler v2.3 (“Newbler”) using default settings, resulting in average read depths of >50× for the mitogenomes of each species. The mitogenome of *A. githago* was then manually assembled into a master circle conformation based on the Newbler assembly graph as described previously (Sloan, Alverson, Wu, et al. 2012). We did not attempt to manually close the assembly of the larger and more complex *S. chalcedonica* mitogenome. In order to correct the known high insertion and deletion error rates associated with long homopolymer regions from 454 sequencing, we performed Illumina sequencing on total-cellular DNA from an individual from the same *A. githago* maternal family (Giles County) used for mitogenome sequencing. DNA was extracted from a 3-day old seedling using a CTAB protocol (Doyle and Doyle 1987). An Illumina library was constructed with the NEBNext Ultra II FS DNA Library Prep Kit, using approximately 30 ng of input DNA, a 20 min fragmentation time, and 8 cycles of PCR amplification. The library was dual-indexed and multiplexed with other libraries on a NovaSeq 6000 S4 sequencing lane (paired-end, 150-bp reads) at the University of Colorado Cancer Center, generating 49.6M read pairs. These reads were then mapped to the mitogenome to correct homopolymer length errors in an iterative fashion as described previously (Sloan, Alverson, Wu, et al. 2012).

The *A. githago* mitogenome was annotated with Mitofy (Alverson, et al. 2010), BLAST homology searches, and tRNAscan-SE (Chan and Lowe 2019). tRNAscan-SE was also used to identify tRNA genes in the mitochondrial contigs of the *S. chalcedonica* assembly. The mitogenome plot (Supp. Fig. 1) was produced with OGDRAW version 1.3.1 (Greiner, et al. 2019). The annotated *A. githago* mitogenome sequence and the contigs from the *S. chalcedonica* assembly were deposited to GenBank (accession numbers MW553037 and MW580967-MW581000, respectively). The raw 454 and Illumina reads are available via NCBI SRA (accessions SRX9983705, SRX9983703, and SRX9983702).

### Tissue growth for tRNA analysis

*Agrostemma githago* seeds were obtained from the Kew Gardens Millennium Seed Bank (Kew 0053084), germinated in small plastic pots with Pro-Mix BX General Purpose soil supplemented with vermiculite and perlite, and grown in growth chambers held at 23°C with a 16-hr/8-hr light/dark cycle (light intensity of 100 μE·m^−2^·s^−1^). *Silene latifolia* seeds were from a female derived from the line originally used for mitogenome sequencing (Sloan, et al. 2010) fertilized by a male obtained from the Kew Gardens Millennium Seed Bank (Kew 32982). Leaf tissue for *S. tuberosum* was generated by planting seed potato (tubers) cultivar White Kennebec. *Silene latifolia*, *S. noctiflora* BRP (Wu, et al. 2015), and *S. vulgaris* S9L (Sloan, Muller, et al. 2012) seeds were germinated in SC7 Cone-tainers (Stuewe and Sons) with the same soil as above on a mist bench under supplemental lighting (16-hr/8-hr light/dark cycle) in the Colorado State University greenhouse. After germination, *S. latifolia*, *S. noctiflora*, and *S. vulgaris* seedlings were moved to a bench with the same 16-hr/8-hr light/dark cycle supplemental lighting until harvest. *S. conica* ABR (Sloan, Alverson, Chuckalovcak, et al. 2012) seeds were germinated in the same soil and containers but grown in a growth room under short-day conditions (10-hr/14-hr light/dark cycle) in order to increase rosette growth. And finally, *S. tuberosum* seed potato was planted in the same soil but in one-gallon pots in the same chamber and short-day conditions (10-hr/14-hr light/dark cycle) to promote leaf tissue growth.

### Mitochondrial isolations

The age of the plants at harvest time ranged depending on species, as certain species required more growth time to produce a sufficient amount of tissue for mitochondrial isolations. *Agrostemma githago* plants were harvested at 4 weeks, *S. conica* at 14 weeks, *S. latifolia* at 6 weeks, *S. noctiflora* at 14 weeks, *S. vulgaris* at 8 months, and *S. tuberosum* at 4 weeks. 75 g of leaf and stem tissue (entire above ground tissue) was collected for each replicate for each respective species. This represented 5 potato plants and anywhere from 24 to 65 *Sileneae* plants per replicate. Leaf tissue was disrupted in a Nutri Ninja Blender for 2 × 2-sec short bursts, and 1 × 4-sec blending in 350 mL of a homogenization buffer containing: 0.3 M sucrose, 5 mM tetrasodium pyrophosphate, 2 mM EDTA, 10 mM KH_2_PO_4_, 1% PVP-40, 1% BSA, 20 mM ascorbic acid, and 5 mM cysteine (pH 7.5-KOH) in a cold room. Homogenate was then poured over four layers of cheese cloth and 1 layer of miracloth and squeezed to filter out all solids. The liquid homogenate was then subjected to differential centrifugation to remove nuclei, plastids, and cellular debris in a Beckman-Coulter Avanti JXN-30 centrifuge with a JA14.50 fixed-angle rotor at 4°C. Homogenate was spun at 500 rcf, 1500 rcf, and 3000 rcf for 10 min at each speed. After each centrifugation step, the supernatant was transferred into a clean centrifuge bottle. Mitochondria were then pelleted by centrifugation for 10 min at 20,000 rcf with the brake off. Supernatant was discarded and the pellet was resuspended using a goat-hair paintbrush and 2 mL wash buffer containing 0.3 M sucrose, 10 mM MOPS, 1 mM EGTA (pH 7.2-KOH). Supernatant was then transferred to 32 mL tubes for centrifugation on a swinging bucket rotor (JS-24.38). The homogenate was centrifuged for 5 min at 3000 rcf to pellet residual plastid and nuclear contaminants, and the supernatant was transferred to a clean 32 mL tube. Mitochondria were once again pelleted by centrifugation for 10 min at 20,000 rcf. The pellet was resuspended with 500 μL wash buffer and a paint brush. The mitochondrial suspension was then added to a glass Dounce homogenizer and homogenized with three strokes.

Homogenized mitochondria were then suspended on top of a Percoll gradient with the following Percoll density layers, 18%, 25%, 50%. The gradient was then centrifuged at 40,000 rcf for 45 min with the brake off. The mitochondrial band at the 25%:50% interface was then aspirated off of the gradient and diluted with 30 mL of wash buffer. The diluted mitochondria were then centrifuged at 20,000 rcf for 10 min. The supernatant was vacuum aspirated, and the mitochondrial pellet was resuspended in a fresh 30 mL of wash buffer and centrifuged at 10,000 rcf for 10 min. The supernatant was vacuum aspirated, and the mitochondrial pellet was resuspended in 1000 μL of fresh wash buffer. Resuspended mitochondria were centrifuged at 10,000 rcf for 10 min. Supernatant was removed with a pipette and the mitochondrial pellet immediately went into RNA extraction procedures using 1000 μL TRIzol following the manufacturer’s RNA extraction protocol.

The total-cellular samples for *A. githago, S. latifolia, and S. noctiflora* were produced by freezing an entire, single plant excluding any root tissue in liquid nitrogen and grinding with a mortar and pestle into powder and performing an RNA extraction using the Trizol manufacturer’s protocol. In the case of *S. tuberosum*, *S. vulgaris, and S. conica*, one shoot from one plant with all leaves was used.

Only two replicates of *S. latifolia* were performed because of tissue limitations, and only two mitochondrial isolations from *S. vulgaris* were used in downstream analysis because of an error quantifying the AlkB/RNA ratio in one of the isolations. Two replicates of mitochondrial isolations and total cellular samples were produced for *S. tuberosum.* Three replicates (mitochondrial isolations and total-cellular samples) were produced for all other species.

### AlkB treatment and YAMAT-seq library construction

See Warren, et al. (2021) for detailed protocols for AlkB expression/purification, RNA treatment and YAMAT-seq (tRNA-seq) library construction. Briefly, a plasmid containing cloned wild type AlkB protein (pET24a-AlkB deltaN11 [plasmid #73622]) was obtained from Addgene (http://www.addgene.org/), and the AlkB protein was expressed and purified at the CSU Biochemistry and Molecular Biology Protein Expression and Purification Facility.

For the demethylation (AlkB) reaction, 6 μg of total RNA was treated with 250 pmols of AlkB for 1 hr at 37°C in a reaction buffer containing 70 μM ammonium iron(II) sulfate hexahydrate, 0.93 mM α-ketoglutaric acid disodium salt dihydrate, 1.86 mM ascorbic acid, and 46.5 mM HEPES (pH 8.0-HCl). The reaction was quenched by adding 4 μl of 100 mM EDTA. RNA was then extracted with a phenol-chloroform RNA extraction, followed by an ethanol precipitation with the addition of 0.08 μg of RNase-free glycogen and resuspension in RNase-free water. RNA integrity was checked on a TapeStation 2200.

To remove amino acids attached to the 3’-end of tRNAs prior to adapter ligation, the demethylated RNA was deacylated in 20 mM Tris HCl (pH 9.0) at 37°C for 40 min. Following deacylation, adapter ligation was performed using a modified protocol from Shigematsu, et al. (2017). A 9 μl reaction volume containing 1 μg of deacylated RNA and 1 pmol of each Y-5’ adapter (4 pmols total) and 4 pmols of the Y-3’ adapter was incubated at 90°C for 2 min. 1 μl of an annealing buffer containing 500 mM Tris-HCl (pH 8.0) and 100 mM MgCl2 was added to the reaction mixture and incubated for 15 min at 37 °C. Ligation was performed by adding 1 unit of T4 RNA Ligase 2 enzyme (New England Biolabs) in 10 μl of 1X reaction buffer and incubating the reaction at 37°C for 60 min, followed by overnight incubation at 4°C. Reverse transcription of ligated RNA was performed using SuperScript IV (Invitrogen) according to the manufacturer’s protocol.

The resulting cDNA was then amplified by PCR using NEBNext PCR Master mix with 7 μl of the reverse transcription reaction. Ten cycles of PCR were performed for *A. githago*, *S. noctiflora,* and *S. vulgaris*. To generate greater library yields, we increased the number of PCR cycles to 12 cycles for *S. conica*, *S. latifolia*, and *S. tuberosum.* PCR was performed on a on a Bio-Rad C1000 Touch thermal cycler with an initial 1 min incubation at 98°C and 10/12 cycles of 30 s at 98°C, 30 s at 60°C and 30 s at 72°C, followed by 5 min at 72°C. Size selection of the resulting PCR products was done on a BluePippin (Sage Science) with Q3 3% agarose gel cassettes following the manufacturer’s protocol. The size selection parameters were set to a range of 180-231 bp. Size-selected products were then cleaned using solid phase reversable immobilization (SPRI) beads and resuspended in 10 mM Tris (pH 8.0). Libraries were dual-indexed and sequenced on an Illumina NovaSeq 6000 S4 lane with paired-end, 150-bp reads at the University of Colorado Cancer Center. YAMAT-seq reads for *A. githago*, *S. conica*, *S. latifolia*, *S. noctiflora* and *S. tuberosum* are available via the NCBI Sequence Read Archive under BioProject PRJNA698234 and sequencing reads for *S. vulgaris* are available under BioProject PRJNA662108.

### YAMAT-seq read processing

Adapters were trimmed using Cutadapt version 1.16 (Martin 2011) with a nextseq trim quality-cutoff parameter of 20 (option: --nextseq-trim=20). A minimum length filter of 50 bp and a maximum of 95 bp was applied to only retain full-length tRNA sequences and remove a substantial number of reads containing only adapter sequence without an insert (Shigematsu, et al. 2017). BBMerge from the BBTools software package was used to merge R1 and R2 read pairs into a consensus sequence (Bushnell, et al. 2017). Identical reads were summed and collapsed into read families using the FASTQ/A Collapser tool from the FASTX-Toolkit version 0.0.13 (http://hannonlab.cshl.edu/fastx_toolkit/index.html).

### Nuclear genome assemblies and reference tRNA genes

In order to generate reference tRNA gene sequences for each species, nuclear genome assemblies were either generated from Illumina short-read datasets or obtained from published sources. Illumina reads for an *A. githago* nuclear assembly were generated from total-cellular DNA extracted from a 5-month-old plant germinated from a line started from Kew 0053084 seeds. DNA was extracted using a Qiagen DNeasy kit, and the NEBNext Ultra II FS DNA Library Prep Kit was used as described above. Sequencing was performed on an Illumina NovaSeq 6000 S4 Lane with paired-end 150-bp reads at the University of Colorado Cancer Center. Shotgun sequencing reads for *A. githago* are available via the NCBI Sequence Read Archive under BioProject PRJNA698248. Illumina reads for an *S. vulgaris* were generated from plants were grown from seeds collected from natural populations in Seefield, Austria. Leaf tissue was collected from a single hermaphrodite plant that had been crossed to the F2 generation and grown under standard greenhouse conditions at the University of Virginia. Genomic DNA was extracted with a modified CTAB protocol (Doyle and Doyle 1987) and sent to Global Biologics (Columbia, MO) for the construction of Illumina libraries with 180 bp inserts. Paired-end (2 x100) reads were sequenced at the Yale Center for Genome Analysis on a single Illumina HiSeq 2000 lane. Raw reads have been deposited into the NCBI Short Read Archive (SRA) with accession SRX10073976. *Silene conica* ABR Illumina reads were obtained from a previously published dataset (Broz, et al. 2021).

Genomic reads for *A. githago, S. conica, S. vulgaris* were trimmed with cutadapt ver. 1.16 (Martin 2011) with a quality-cutoff parameter of 10 for the 3’ end of each read and a minimum length filter of 5 bp. Reads were then assembled using SPAdes v3.11.0, (Nurk, et al. 2013) using the following command options: -k 21,33,55,77,99 -t 40 -m 900. Assemblies for *S. noctiflora* OPL (GenBank assembly accession: VHZZ00000000.1) and *S. latifolia* (GenBank assembly accession: QBIE00000000.1) were taken from previous studies (Krasovec, et al. 2018; Williams, et al. 2020).

A database of nuclear tRNA reference genes was produced by searching each of the nuclear assemblies from *A. githago*, *S. conica, S. latifolia*, *S. noctiflora*, and *S. vulgaris* with tRNAscan-SE. ver. 2.0.3 (Chan and Lowe 2019) using the General search option (-G). Introns predicted from tRNAscan-SE were removed, and identical tRNA sequences were collapsed into a single reference using custom Perl scripts. It is important to note that plant genomes have multiple copies of identical tRNA genes, but for mapping purposes all of these identical sequences are collapsed into a single reference. The nuclear references for *S. tuberosum* were obtained from the curated PlantRNA website (http://seve.ibmp.unistra.fr/plantrna/, downloaded Oct. 26, 2020).

### Mitochondrial genome mapping and stem-loop references

To search for previously undetected mitochondrial tRNA sequences captured by YAMAT-seq, as well as to identify any expressed stem-loop regions modified with a CCA-tail, all YAMAT reads for each species were mapped to the entire mitogenome using BLAST (blastn, e-value of < 1e-6, low complexity regions not filtered [-dust no]). To ensure only high confidence mitogenome hits, reads mapping to the mitogenome were then filtered so that only reads with greater than or equal to 90% identical matches and hit coverage to a mitogenome location were retained. The “closest” function in bedtools ver. 2.27.1 (Quinlan and Hall 2010) was used to assign the closest annotated gene for each of these retained reads, including any reads that mapped/overlapped with a previously annotated tRNA. All reads that did not map to a tRNA gene were then further analyzed to determine if they occurred at the boundary of protein-coding or RNA gene (indicating that these transcripts may be t-elements). Those that did not occur at a gene boundary were considered to represent a previously undocumented stem-loop or tRNA-like structure. Highly expressed t-elements and stem-loops were added to the mapping database as reference sequences.

### Folding predictions of mitochondrial t-elements and stem-loops

Folding prediction analysis for Supp. Fig. 2 was done with RNAFold (ver. 2.4.11, (Lorenz, et al. 2011)), using the using maximum free energy (MFE) model. Because stem-loops also had RT misincorporations at some sites, the YAMAT sequence with 100% nucleotide identity to the mitogenome was used for the folding prediction. Folding diagrams were created with VARNA (ver. 3.9, (Darty, et al. 2009)).

### Contamination removal

Although the vast majority of the YAMAT reads originated from the sampled plant species, there were numerous reads of contaminating origins such as soil bacteria and plant pests. In order to remove reads that did not originate from the plant itself, all unique sequences with three or more reads were BLASTED to the NCBI nucleotide collection (nt) database (posting date of Oct 27, 2019, downloaded Feb 02, 2020) (blastn, e-value of < 1e-3), and the taxonomy information for the top two hits for each read was pulled using the NCBI taxonomy database (downloaded Feb 02, 2020). All queries that did not have the taxon name Viridiplantae in the first two hits were retained to make a contamination database. GenBank files were downloaded for all contaminating accessions that had more than 50 reads assigned, and tRNA annotations were extracted from those GenBank files. Some accessions did not have annotated tRNA genes, so the contaminating hit was added to the database manually. Many of the contaminating references were identical or nearly identical to each other (e.g., many bacterial tRNA sequences were identical or only differed by slight length variation due to annotations). Thus, the contaminant database was collapsed to retain only unique sequences and only the longest reference of otherwise identical sequences. Contaminating sequences were added to the mapping database as reference sequences. We also filtered YAMAT-seq reads that did not map to any reference with at least 90% sequence identity and 80% coverage (see below) to ensure that that any low-abundance, contaminating reads were not retained in the analysis.

### Read mapping

Mapping of processed reads was performed with a custom Perl script previously published in Warren, et al. (2021) that can be found at (https://github.com/warrenjessica/YAMAT-scripts). Briefly, each read was BLASTed (blastn, e-value threshold of < 1e-6) to all references. A mapped read was retained if it had 90% or more sequence identity and at least 80% coverage to a reference sequence. This ensured that full-length tRNA sequences were used in analysis but also allowed for some mismatches due to reverse transcription-induced misincorporations (Warren, et al. 2021). Furthermore, only sequences that uniquely mapped to a single reference were retained for downstream analysis. All mapped and filtered reads were then counted for differential expression analysis.

### EdgeR differential expression analysis

Differential expression analysis was done on mapped counts with the R package EdgeR version 3.24.3 (Robinson, et al. 2010) using a gene-wise negative binomial generalized linear models with quasi-likelihood tests (glmQLFit). Library sizes were normalized with the function calcNormFactors(y) and low-abundance genes were filtered by expression using the filterByExpr(y) function. Because misincorporations in cDNA are common when reverse transcribing tRNAs, lowly detected (< 2 counts per million or CPM) sequences in this analysis likely represent rare misincorporation profiles of tRNA sequences (and not true unique genes) which occasionally mapped to unique references. To summarize the level of mitochondrial enrichment by anticodon (isodecoder) family, a weighted average enrichment level for all tRNAs with the same anticodon was calculated and presented in Figures 6 and 7. The average enrichment was calculating by weighting each sequence by its percentage of total expression (CPM) within the isodecoder family. Count data as well as differential expression results can be found in Supp. Tables 2-8, and the weighted enrichment for each anticodon group for each species can be found in Supp. Table 11.

## Supporting information

All Supplemental Figures

## Acknowledgements

We would like to thank Laura Bergner for mitochondrial DNA extractions of *Agrostemma githago* and *Silene chalcedonica* and Zhiqiang Wu for generation of preliminary tRNA-seq data. This work was supported by an NSF award (MCB-2048407), a Chateaubriand Fellowship from the Embassy of France in the United States, an NSF GAUSSI graduate research fellowship (DGE-1450032), and by the Centre National de la Recherche Scientifique (CNRS) and the “Initiative d’Excellence” (MITOCROSS ANR-11-LABX-0057).

## References

Alverson AJ, Wei XX, Rice DW, Stern DB, Barry K, Palmer JD. 2010. Insights into the Evolution of Mitochondrial Genome Size from Complete Sequences of Citrullus lanatus and Cucurbita pepo (Cucurbitaceae). Molecular Biology and Evolution 27:1436–1448.

Anderson S, Bankier AT, Barrell BG, de Bruijn MH, Coulson AR, Drouin J, Eperon IC, Nierlich DP, Roe BA, Sanger F, et al. 1981. Sequence and organization of the human mitochondrial genome. Nature 290:457–465.

Ayan GB, Park HJ, Gallie J. 2020. The birth of a bacterial tRNA gene by large-scale, tandem duplication events. Elife 9.

Binder S, Marchfelder A, Brennicke A. 1994. RNA editing of tRNA(Phe) and tRNA(Cys) in mitochondria of Oenothera berteriana is initiated in precursor molecules. Mol Gen Genet 244:67–74.

Binder S, Marchfelder A, Brennicke A, Wissinger B. 1992. RNA editing in trans-splicing intron sequences of nad2 mRNAs in Oenothera mitochondria. J Biol Chem 267:7615–7623.

Boore JL, Macey JR, Medina M. 2005. Sequencing and comparing whole mitochondrial genomes of animals. Methods Enzymol 395:311–348.

Brandvain Y, Wade MJ. 2009. The functional transfer of genes from the mitochondria to the nucleus: the effects of selection, mutation, population size and rate of self-fertilization. Genetics 182:1129–1139.

Brown WM, George M, Jr., Wilson AC. 1979. Rapid evolution of animal mitochondrial DNA. Proc Natl Acad Sci U S A 76:1967–1971.

Broz AK, Waneka G, Wu Z, Fernandes Gyorfy M, Sloan DB. 2021. Detecting de novo mitochondrial mutations in angiosperms with highly divergent evolutionary rates. Genetics 218.

Brubacher-Kauffmann S, Marechal-Drouard L, Cosset A, Dietrich A, Duchêne AM. 1999. Differential import of nuclear-encoded tRNAGly isoacceptors into solanum Tuberosum mitochondria. Nucleic Acids Res 27:2037–2042.

Bushnell B, Rood J, Singer E. 2017. BBMerge - Accurate paired shotgun read merging via overlap. Plos One 12.

Calvo SE, Mootha VK. 2010. The mitochondrial proteome and human disease. Annu Rev Genomics Hum Genet 11:25–44.

Ceci LR, Veronico P, Siculella L, Gallerani R. 1995. Identification and mapping of trnI, trnE and trnfM genes in the sunflower mitochondrial genome. DNA Seq 5:315–318.

Chan PP, Lowe TM. 2019. tRNAscan-SE: Searching for tRNA Genes in Genomic Sequences. Methods Mol Biol 1962:1–14.

Chapdelaine Y, Bonen L. 1991. The wheat mitochondrial gene for subunit I of the NADH dehydrogenase complex: a trans-splicing model for this gene-in-pieces. Cell 65:465–472.

Chimnaronk S, Gravers Jeppesen M, Suzuki T, Nyborg J, Watanabe K. 2005. Dual-mode recognition of noncanonical tRNAs(Ser) by seryl-tRNA synthetase in mammalian mitochondria. EMBO J 24:3369–3379.

Clary DO, Wolstenholme DR. 1984. The Drosophila mitochondrial genome. Oxf Surv Eukaryot Genes 1:1–35.

Cozen AE, Quartley E, Holmes AD, Hrabeta-Robinson E, Phizicky EM, Lowe TM. 2015. ARM-seq: AlkB-facilitated RNA methylation sequencing reveals a complex landscape of modified tRNA fragments. Nature Methods 12:879-+.

Darty K, Denise A, Ponty Y. 2009. VARNA: Interactive drawing and editing of the RNA secondary structure. Bioinformatics 25:1974–1975.

Delage L, Dietrich A, Cosset A, Marechal-Drouard L. 2003. In vitro import of a nuclearly encoded tRNA into mitochondria of Solanum tuberosum. Mol Cell Biol 23:4000–4012.

Dewe JM, Whipple JM, Chernyakov I, Jaramillo LN, Phizicky EM. 2012. The yeast rapid tRNA decay pathway competes with elongation factor 1A for substrate tRNAs and acts on tRNAs lacking one or more of several modifications. RNA 18:1886–1896.

Doyle JJ, Doyle JL. 1987. A rapid DNA isolation procedure for small quantities of fresh leaf tissue. PHYTOCHEMICAL BULLETIN v.19(1):11–15.

Duchêne AM, Marechal-Drouard L. 2001. The chloroplast-derived trnW and trnM-e genes are not expressed in Arabidopsis mitochondria. Biochem Biophys Res Commun 285:1213–1216.

Duchêne AM, Pujol C, Marechal-Drouard L. 2009. Import of tRNAs and aminoacyl-tRNA synthetases into mitochondria. Curr Genet 55:1–18.

Forner J, Weber B, Thuss S, Wildum S, Binder S. 2007. Mapping of mitochondrial mRNA termini in Arabidopsis thaliana: t-elements contribute to 5’ and 3’ end formation. Nucleic Acids Res 35:3676–3692.

Giegé R, Sissler M, Florentz C. 1998. Universal rules and idiosyncratic features in tRNA identity. Nucleic Acids Research 26:5017–5035.

Glover KE, Spencer DF, Gray MW. 2001. Identification and structural characterization of nucleus-encoded transfer RNAs imported into wheat mitochondria. J Biol Chem 276:639–648.

Gray MW. 2012. Mitochondrial evolution. Cold Spring Harb Perspect Biol 4:a011403.

Greiner S, Lehwark P, Bock R. 2019. OrganellarGenomeDRAW (OGDRAW) version 1.3.1: expanded toolkit for the graphical visualization of organellar genomes. Nucleic Acids Res 47:W59–W64.

Grosjean H, Westhof E. 2016. An integrated, structure- and energy-based view of the genetic code. Nucleic Acids Res 44:8020–8040.

Guo W, Zhu A, Fan W, Adams RP, Mower JP. 2020. Extensive Shifts from Cis- to Trans-splicing of Gymnosperm Mitochondrial Introns. Molecular Biology and Evolution 37:1615–1620.

Haen KM, Pett W, Lavrov DV. 2010. Parallel Loss of Nuclear-Encoded Mitochondrial Aminoacyl-tRNA Synthetases and mtDNA-Encoded tRNAs in Cnidaria. Molecular Biology and Evolution 27:2216–2219.

Hancock K, Hajduk SL. 1990. The mitochondrial tRNAs of Trypanosoma brucei are nuclear encoded. J Biol Chem 265:19208–19215.

Huynen MA, Duarte I, Szklarczyk R. 2013. Loss, replacement and gain of proteins at the origin of the mitochondria. Biochimica Et Biophysica Acta-Bioenergetics 1827:224–231.

Iams KP, Heckman JE, Sinclair JH. 1985. Sequence of histidyl tRNA, present as a chloroplast insert in mtDNA ofZea mays. Plant Molecular Biology 4:225–232.

Jafari F, Zarre S, Gholipour A, Eggens F, Rabeler RK, Oxelman B. 2020. A new taxonomic backbone for the infrageneric classification of the species-rich genus Silene (Caryophyllaceae). Taxon 69:337–368.

Juhling F, Morl M, Hartmann RK, Sprinzl M, Stadler PF, Putz J. 2009. tRNAdb 2009: compilation of tRNA sequences and tRNA genes. Nucleic Acids Res 37:D159–162.

Juhling T, Duchardt-Ferner E, Bonin S, Wohnert J, Putz J, Florentz C, Betat H, Sauter C, Morl M. 2018. Small but large enough: structural properties of armless mitochondrial tRNAs from the nematode Romanomermis culicivorax. Nucleic Acids Res 46:9170–9180.

Kitazaki K, Kubo T, Kagami H, Matsumoto T, Fujita A, Matsuhira H, Matsunaga M, Mikami T. 2011. A horizontally transferred tRNA(Cys) gene in the sugar beet mitochondrial genome: evidence that the gene is present in diverse angiosperms and its transcript is aminoacylated. Plant J 68:262–272.

Knie N, Polsakiewicz M, Knoop V. 2015. Horizontal Gene Transfer of Chlamydial-Like tRNA Genes into Early Vascular Plant Mitochondria. Molecular Biology and Evolution 32:629–634.

Knoop V, Schuster W, Wissinger B, Brennicke A. 1991. Trans splicing integrates an exon of 22 nucleotides into the nad5 mRNA in higher plant mitochondria. EMBO J 10:3483–3493.

Krasovec M, Chester M, Ridout K, Filatov DA. 2018. The Mutation Rate and the Age of the Sex Chromosomes in Silene latifolia. Curr Biol 28:1832–1838 e1834.

Kubo T, Nishizawa S, Sugawara A, Itchoda N, Estiati A, Mikami T. 2000. The complete nucleotide sequence of the mitochondrial genome of sugar beet (Beta vulgaris L.) reveals a novel gene for tRNA(Cys)(GCA). Nucleic Acids Res 28:2571–2576.

Kumar R, Marechal-Drouard L, Akama K, Small I. 1996. Striking differences in mitochondrial tRNA import between different plant species. Mol Gen Genet 252:404–411.

Lavrov DV, Pett W. 2016. Animal Mitochondrial DNA as We Do Not Know It: mt-Genome Organization and Evolution in Nonbilaterian Lineages. Genome Biol Evol 8:2896–2913.

Lorenz R, Bernhart SH, Honer Zu Siederdissen C, Tafer H, Flamm C, Stadler PF, Hofacker IL. 2011. ViennaRNA Package 2.0. Algorithms Mol Biol 6:26.

Magallon S, Gomez-Acevedo S, Sanchez-Reyes LL, Hernandez-Hernandez T. 2015. A metacalibrated time-tree documents the early rise of flowering plant phylogenetic diversity. New Phytologist 207:437–453.

Marechal-Drouard L, Guillemaut P, Cosset A, Arbogast M, Weber F, Weil JH, Dietrich A. 1990. Transfer RNAs of potato (Solanum tuberosum) mitochondria have different genetic origins. Nucleic Acids Res 18:3689–3696.

Marechal-Drouard L, Ramamonjisoa D, Cosset A, Weil JH, Dietrich A. 1993. Editing corrects mispairing in the acceptor stem of bean and potato mitochondrial phenylalanine transfer RNAs. Nucleic Acids Res 21:4909–4914.

Martin M. 2011. Cutadapt removes adapter sequences from high-throughput sequencing reads. EMBnet. journal 17:10–12.

Meiklejohn CD, Holmbeck MA, Siddiq MA, Abt DN, Rand DM, Montooth KL. 2013. An Incompatibility between a Mitochondrial tRNA and Its Nuclear-Encoded tRNA Synthetase Compromises Development and Fitness in Drosophila. Plos Genetics 9.

Mercer TR, Neph S, Dinger ME, Crawford J, Smith MA, Shearwood AM, Haugen E, Bracken CP, Rackham O, Stamatoyannopoulos JA, et al. 2011. The human mitochondrial transcriptome. Cell 146:645–658.

Michaud M, Cognat V, Duchêne AM, Marechal-Drouard L. 2011. A global picture of tRNA genes in plant genomes. Plant J 66:80–93.

Mower JP, Touzet P, Gummow JS, Delph LF, Palmer JD. 2007. Extensive variation in synonymous substitution rates in mitochondrial genes of seed plants. BMC Evol Biol 7:135.

Nurk S, Bankevich A, Antipov D, Gurevich AA, Korobeynikov A, Lapidus A, Prjibelski AD, Pyshkin A, Sirotkin A, Sirotkin Y, et al. 2013. Assembling single-cell genomes and mini-metagenomes from chimeric MDA products. J Comput Biol 20:714–737.

Pawar K, Shigematsu M, Loher P, Honda S, Rigoutsos I, Kirino Y. 2019. Exploration of CCA-added RNAs revealed the expression of mitochondrial non-coding RNAs regulated by CCA-adding enzyme. RNA Biol 16:1817–1825.

Petersen G, Cuenca A, Moller IM, Seberg O. 2015. Massive gene loss in mistletoe (Viscum, Viscaceae) mitochondria. Sci Rep 5:17588.

Peterson KJ, Lyons JB, Nowak KS, Takacs CM, Wargo MJ, McPeek MA. 2004. Estimating metazoan divergence times with a molecular clock. Proc Natl Acad Sci U S A 101:6536–6541.

Pett W, Lavrov DV. 2015. Cytonuclear Interactions in the Evolution of Animal Mitochondrial tRNA Metabolism. Genome Biol Evol 7:2089–2101.

Pons J, Bover P, Bidegaray-Batista L, Arnedo MA. 2019. Arm-less mitochondrial tRNAs conserved for over 30 millions of years in spiders. BMC Genomics 20:665.

Pujol C, Bailly M, Kern D, Marechal-Drouard L, Becker H, Duchêne AM. 2008. Dual-targeted tRNA-dependent amidotransferase ensures both mitochondrial and chloroplastic Gln-tRNAGln synthesis in plants. Proc Natl Acad Sci U S A 105:6481–6485.

Qiu YL, Palmer JD. 2004. Many independent origins of trans splicing of a plant mitochondrial group II intron. J Mol Evol 59:80–89.

Quinlan AR, Hall IM. 2010. BEDTools: a flexible suite of utilities for comparing genomic features. Bioinformatics 26:841–842.

Raina M, Ibba M. 2014. tRNAs as regulators of biological processes. Front Genet 5:171.

Rice DW, Alverson AJ, Richardson AO, Young GJ, Sanchez-Puerta MV, Munzinger J, Barry K, Boore JL, Zhang Y, dePamphilis CW, et al. 2013. Horizontal transfer of entire genomes via mitochondrial fusion in the angiosperm Amborella. Science 342:1468–1473.

Richardson AO, Rice DW, Young GJ, Alverson AJ, Palmer JD. 2013. The “fossilized” mitochondrial genome of Liriodendron tulipifera: ancestral gene content and order, ancestral editing sites, and extraordinarily low mutation rate. BMC Biol 11:29.

Robinson MD, McCarthy DJ, Smyth GK. 2010. edgeR: a Bioconductor package for differential expression analysis of digital gene expression data. Bioinformatics 26:139–140.

Salinas-Giegé T, Giegé R, Giegé P. 2015. tRNA biology in mitochondria. Int J Mol Sci 16:4518–4559.

Sanchez-Puerta MV, Edera A, Gandini CL, Williams AV, Howell KA, Nevill PG, Small I. 2019. Genome-scale transfer of mitochondrial DNA from legume hosts to the holoparasite Lophophytum mirabile (Balanophoraceae). Molecular Phylogenetics and Evolution 132:243–250.

Shigematsu M, Honda S, Loher P, Telonis AG, Rigoutsos I, Kirino Y. 2017. YAMAT-seq: an efficient method for high-throughput sequencing of mature transfer RNAs. Nucleic Acids Research 45.

Simpson AM, Suyama Y, Dewes H, Campbell DA, Simpson L. 1989. Kinetoplastid Mitochondria Contain Functional Transfer-Rnas Which Are Encoded in Nuclear-DNA and Also Contain Small Minicircle and Maxicircle Transcripts of Unknown Function. Nucleic Acids Research 17:5427–5445.

Mitochondrial Aminoacyl-tRNA Synthetases [Internet]. 2005 Release 2000-2013. Landes Bioscience. Available from https://www.ncbi.nlm.nih.gov/books/NBK6033/.

Skippington E, Barkman TJ, Rice DW, Palmer JD. 2015. Miniaturized mitogenome of the parasitic plant Viscum scurruloideum is extremely divergent and dynamic and has lost all nad genes. Proc Natl Acad Sci U S A 112:E3515–3524.

Sloan DB, Alverson AJ, Chuckalovcak JP, Wu M, McCauley DE, Palmer JD, Taylor DR. 2012. Rapid Evolution of Enormous, Multichromosomal Genomes in Flowering Plant Mitochondria with Exceptionally High Mutation Rates. Plos Biology 10.

Sloan DB, Alverson AJ, Storchova H, Palmer JD, Taylor DR. 2010. Extensive loss of translational genes in the structurally dynamic mitochondrial genome of the angiosperm Silene latifolia. BMC Evol Biol 10:274.

Sloan DB, Alverson AJ, Wu M, Palmer JD, Taylor DR. 2012. Recent acceleration of plastid sequence and structural evolution coincides with extreme mitochondrial divergence in the angiosperm genus Silene. Genome Biol Evol 4:294–306.

Sloan DB, Muller K, McCauley DE, Taylor DR, Storchova H. 2012. Intraspecific variation in mitochondrial genome sequence, structure, and gene content in Silene vulgaris, an angiosperm with pervasive cytoplasmic male sterility. New Phytologist 196:1228–1239.

Sloan DB, Oxelman B, Rautenberg A, Taylor DR. 2009. Phylogenetic analysis of mitochondrial substitution rate variation in the angiosperm tribe Sileneae. BMC Evol Biol 9:260.

Sloan DB, Triant DA, Forrester NJ, Bergner LM, Wu M, Taylor DR. 2014. A recurring syndrome of accelerated plastid genome evolution in the angiosperm tribe Sileneae (Caryophyllaceae). Molecular Phylogenetics and Evolution 72:82–89.

Sloan DB, Warren JM, Williams AM, Wu Z, Abdel-Ghany SE, Chicco AJ, Havird JC. 2018. Cytonuclear integration and co-evolution. Nat Rev Genet 19:635–648.

Small I, Akashi K, Chapron A, Dietrich A, Duchêne AM, Lancelin D, Marechal-Drouard L, Menand B, Mireau H, Moudden Y, et al. 1999. The strange evolutionary history of plant mitochondrial tRNAs and their aminoacyl-tRNA synthetases. Journal of Heredity 90:333–337.

Varre JS, D’Agostino N, Touzet P, Gallina S, Tamburino R, Cantarella C, Ubrig E, Cardi T, Drouard L, Gualberto JM, et al. 2019. Complete Sequence, Multichromosomal Architecture and Transcriptome Analysis of the Solanum tuberosum Mitochondrial Genome. Int J Mol Sci 20.

Vinogradova E, Salinas T, Cognat V, Remacle C, Marechal-Drouard L. 2009. Steady-state levels of imported tRNAs in Chlamydomonas mitochondria are correlated with both cytosolic and mitochondrial codon usages. Nucleic Acids Res 37:1521–1528.

Warren JM, Salinas-Giegé T, Hummel G, Coots NL, Svendsen JM, Brown KC, Drouard L, Sloan DB. 2021. Combining tRNA sequencing methods to characterize plant tRNA expression and post-transcriptional modification. RNA Biology 18:64–78.

Warren JM, Sloan DB. 2020a. Hopeful monsters: Unintended sequencing of famously malformed mite mitochondrial tRNAs reveals widespread expression and processing of sense-antisense pairs. bioRxiv:2020.2009.2008.286963.

Warren JM, Sloan DB. 2020b. Interchangeable parts: The evolutionarily dynamic tRNA population in plant mitochondria. Mitochondrion 52:144–156.

Watanabe Y, Suematsu T, Ohtsuki T. 2014. Losing the stem-loop structure from metazoan mitochondrial tRNAs and co-evolution of interacting factors. Front Genet 5:109.

Williams AM, Itgen MW, Broz AK, Carter OG, Sloan DB. 2020. Long-read transcriptome and other genomic resources for the angiosperm *Silene noctiflora*. bioRxiv:2020.2008.2009.243378.

Williams MA, Johzuka Y, Mulligan RM. 2000. Addition of non-genomically encoded nucleotides to the 3’-terminus of maize mitochondrial mRNAs: truncated rps12 mRNAs frequently terminate with CCA. Nucleic Acids Res 28:4444–4451.

Wilusz JE. 2015. Removing roadblocks to deep sequencing of modified RNAs. Nat Methods 12:821–822.

Wissinger B, Schuster W, Brennicke A. 1991. Trans splicing in Oenothera mitochondria: nad1 mRNAs are edited in exon and trans-splicing group II intron sequences. Cell 65:473–482.

Wu Z, Cuthbert JM, Taylor DR, Sloan DB. 2015. The massive mitochondrial genome of the angiosperm Silene noctiflora is evolving by gain or loss of entire chromosomes. Proc Natl Acad Sci U S A 112:10185–10191.

Yu CH, Liao JY, Zhou H, Qu LH. 2008. The rat mitochondrial Ori L encodes a novel small RNA resembling an ancestral tRNA. Biochem Biophys Res Commun 372:634–638.

Zheng G, Qin Y, Clark WC, Dai Q, Yi C, He C, Lambowitz AM, Pan T. 2015. Efficient and quantitative high-throughput tRNA sequencing. Nat Methods 12:835–837.

