## Supplementary material for "Rapid shifts in mitochondrial tRNA import in a plant lineage with extensive mitochondrial tRNA gene loss": All Supplemental Figures

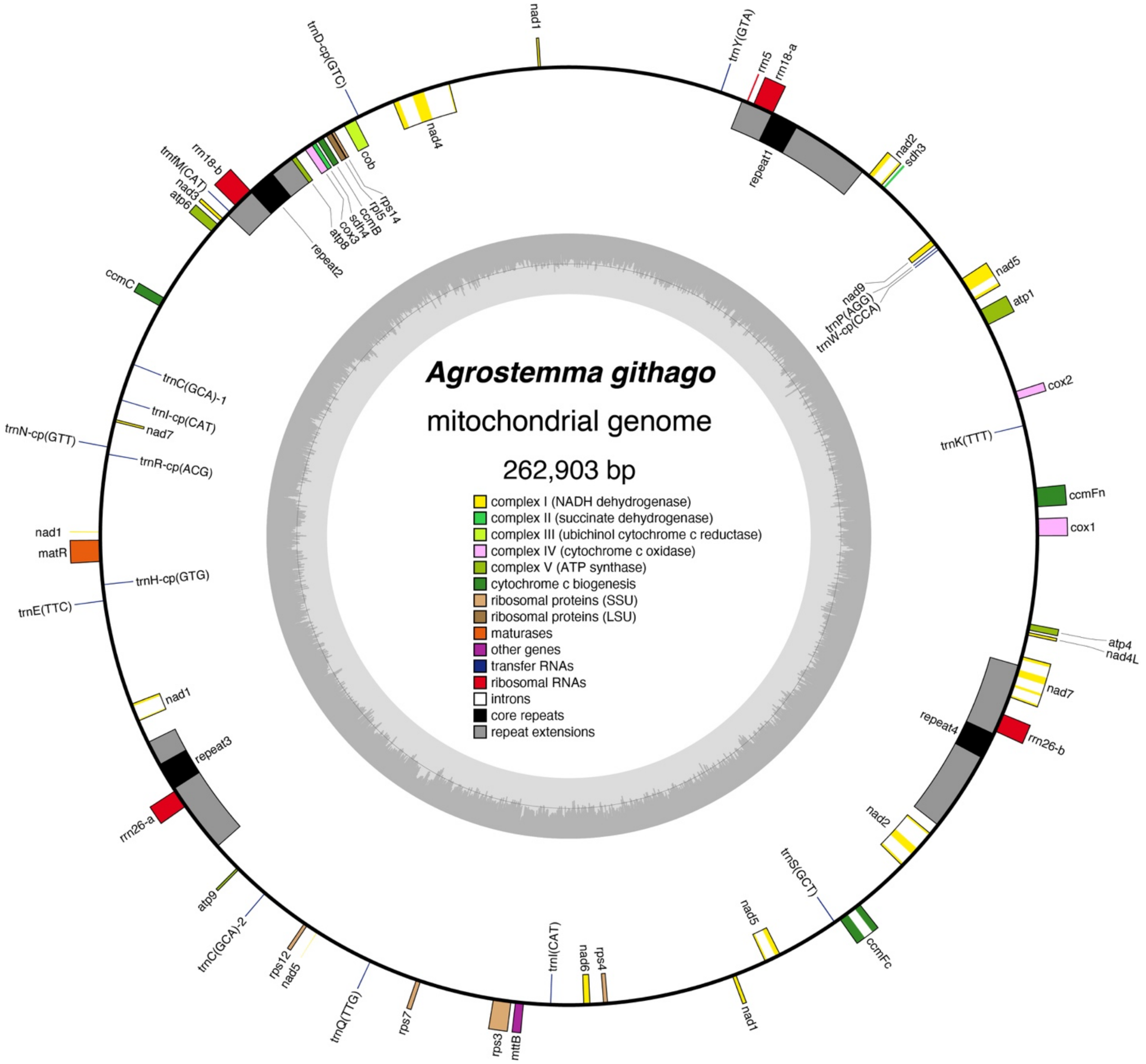


Supp. Fig.1| Mitochondrial genome map of *Agrostemma githago*. This “master circle” represents only one possible configuration of the mitogenome due to multiple recombing repeats. Boxes inside and outside the circle correspond to the clockwise and anticlockwise strands, respectively. The black regions of the repeats designate the identical sequence shared by all four copies. The flanking gray boxes represent extensions to these repeat regions, which are present in two or more (but not all four) copies. The inner gray track represents GC content. Figure was generated with OGDraw v1.3.1(Greiner, et al. 2019)

**T-elements and stem-loops**

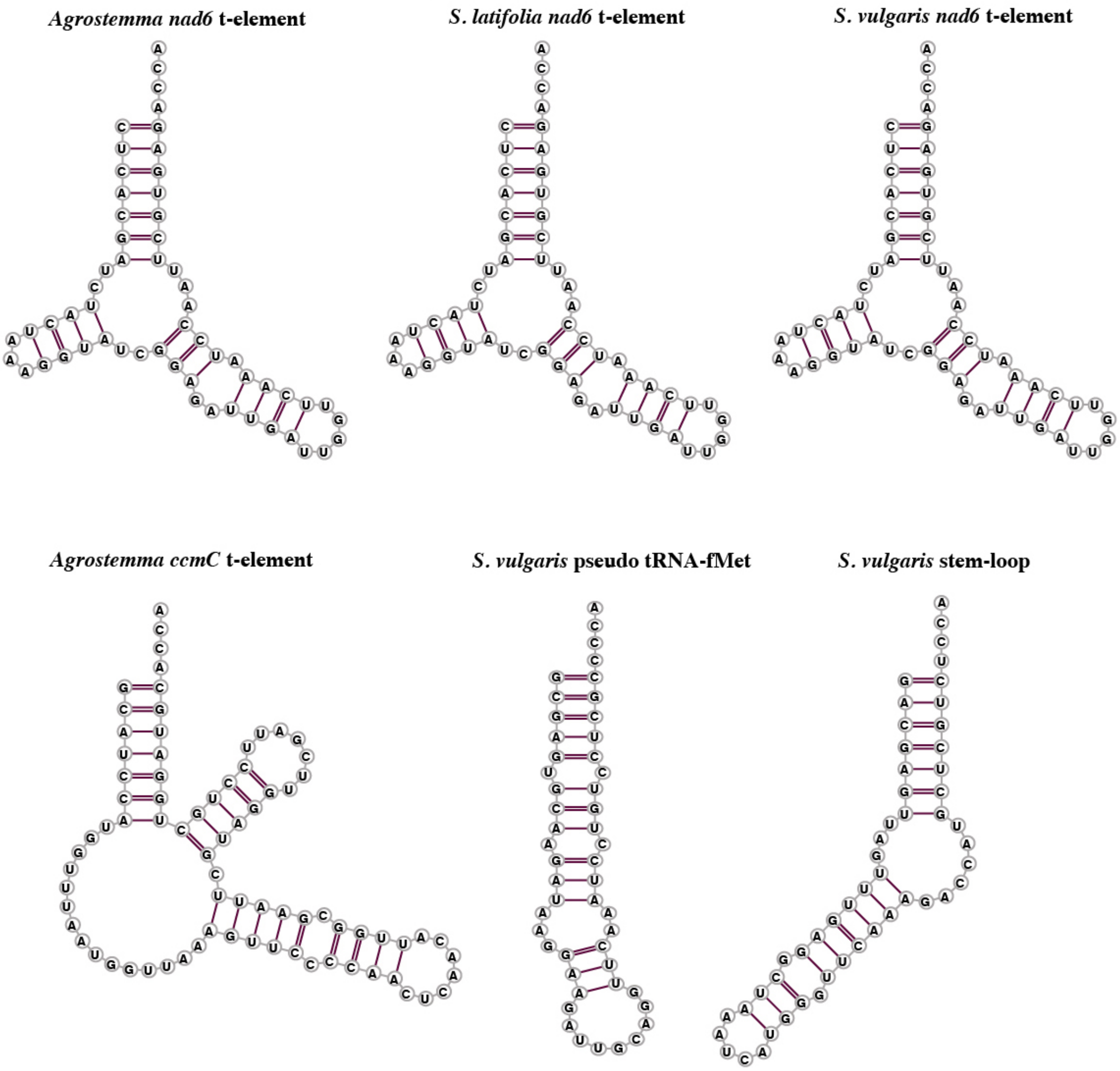


Supp. Fig. 2| Predicted foldings of some of the most highly detected stem-loop and t-element structures. Folding predictions were done using RNAfold (Lorenz, et al. 2011) with a maximum free energy model, and diagrams were created with VARNA (ver. 3.9, (Darty, et al. 2009)).

### Nuclear gene 1009
### alternative folding

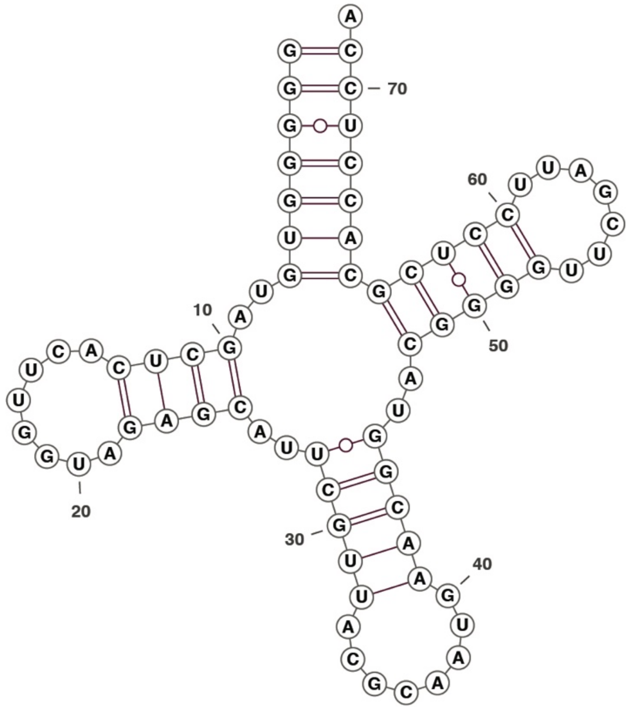


Supp. Fig. 3| Alternative folding prediction of the mitochondrially-enriched, nuclear-encoded tRNA in *S. vulgaris*. Structure presented represents the maximum free energy folding model using the program the program RNAfold (Lorenz, et al. 2011) and diagram was created using the program VARNA ver. 3.9 (Darty, et al. 2009).

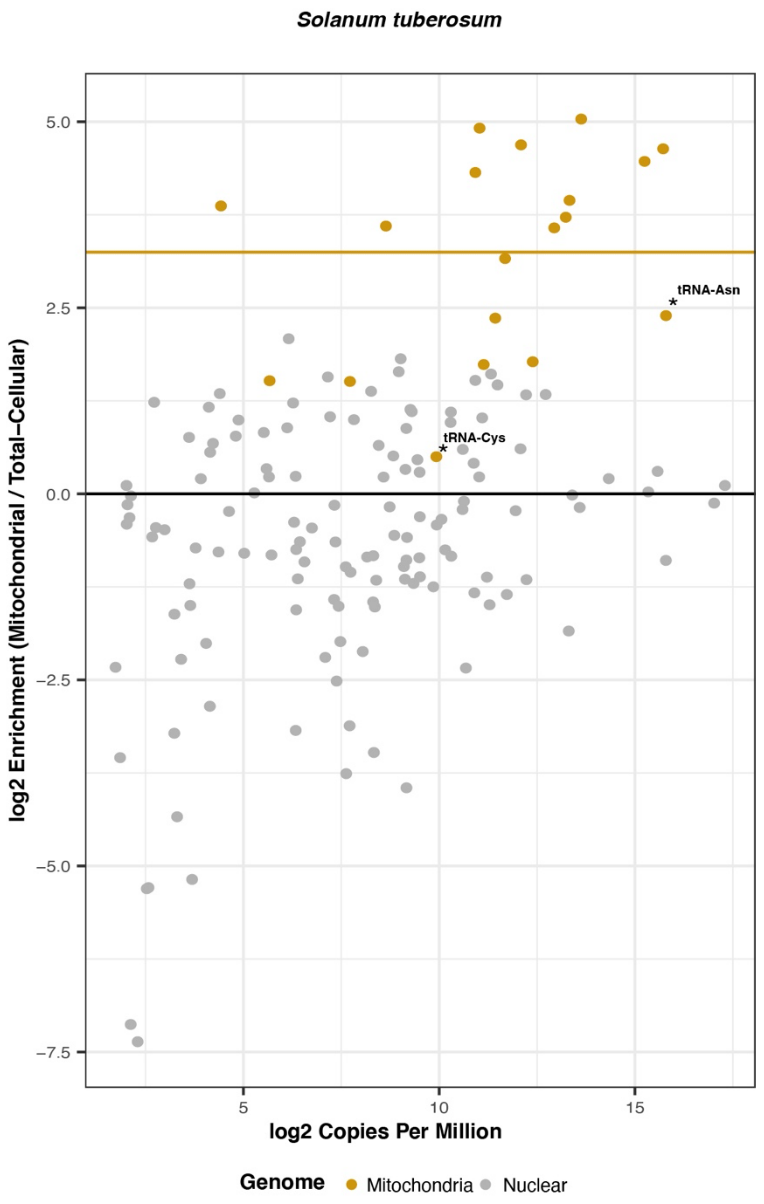


Supp. Fig. 4| Enrichment of mitochondrially encoded and nuclear-encoded tRNAs in the mitochondrial isolate relative to total-cellular sample in *S. tuberosum*. Each dot represents a unique tRNA sequence. The heavy gold line represents the average enrichment of mitochondrial-encoded genes. The genomic origin of the tRNA is indicated by color, with gray being nuclear-encoded and gold being mitochondrial. The two labeled mt-tRNAs (Cys and Asn) are identical in sequence to the plastid-encoded tRNA counterparts.

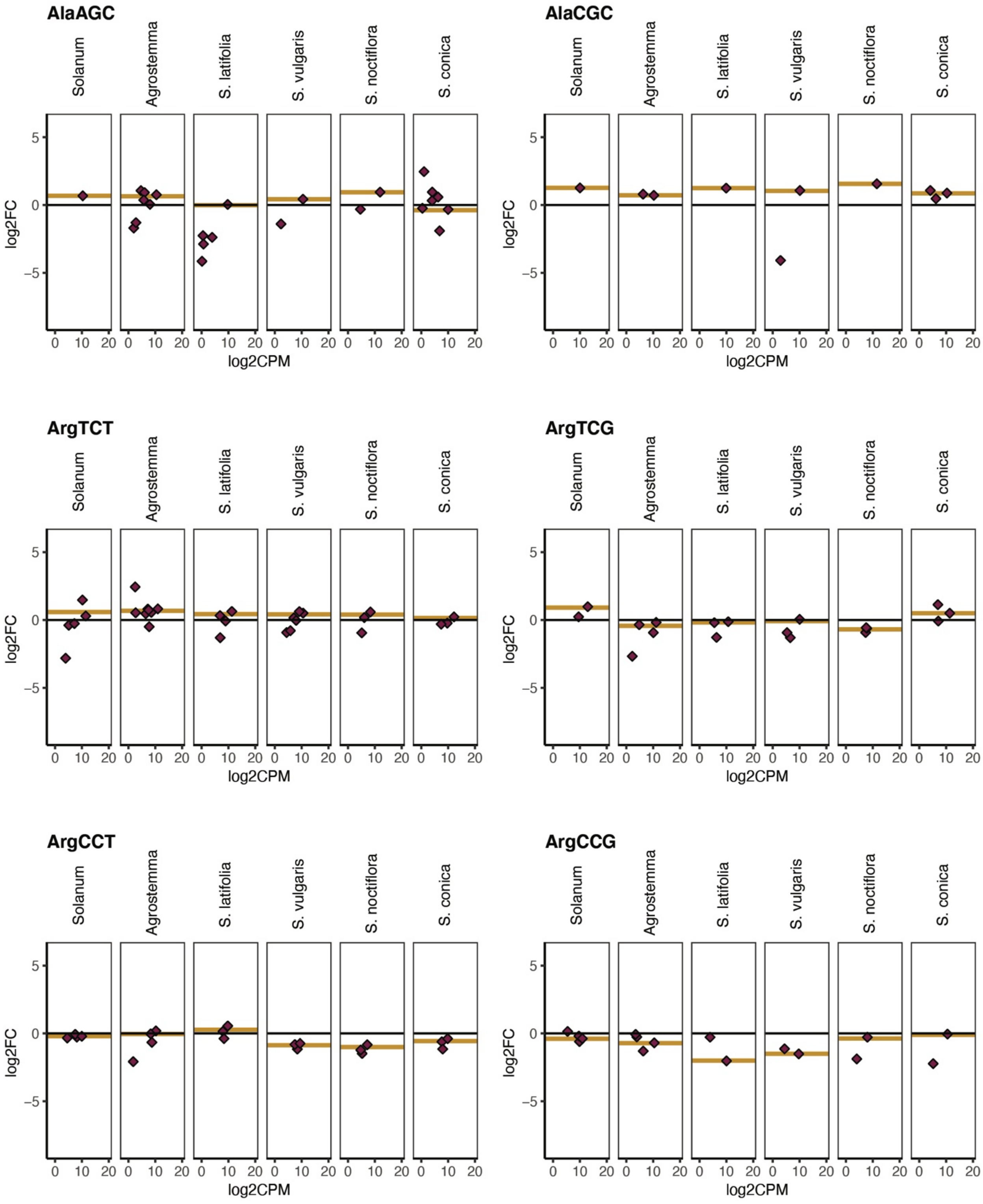

AlaAGC
AlaCGC
ArgTCT
ArgTCG
ArgCCT
ArgCCG
Solanum
Agrostemma
S. latifolia
S. vulgaris
S. noctiflora
S. conica
log2FC
log2CPM

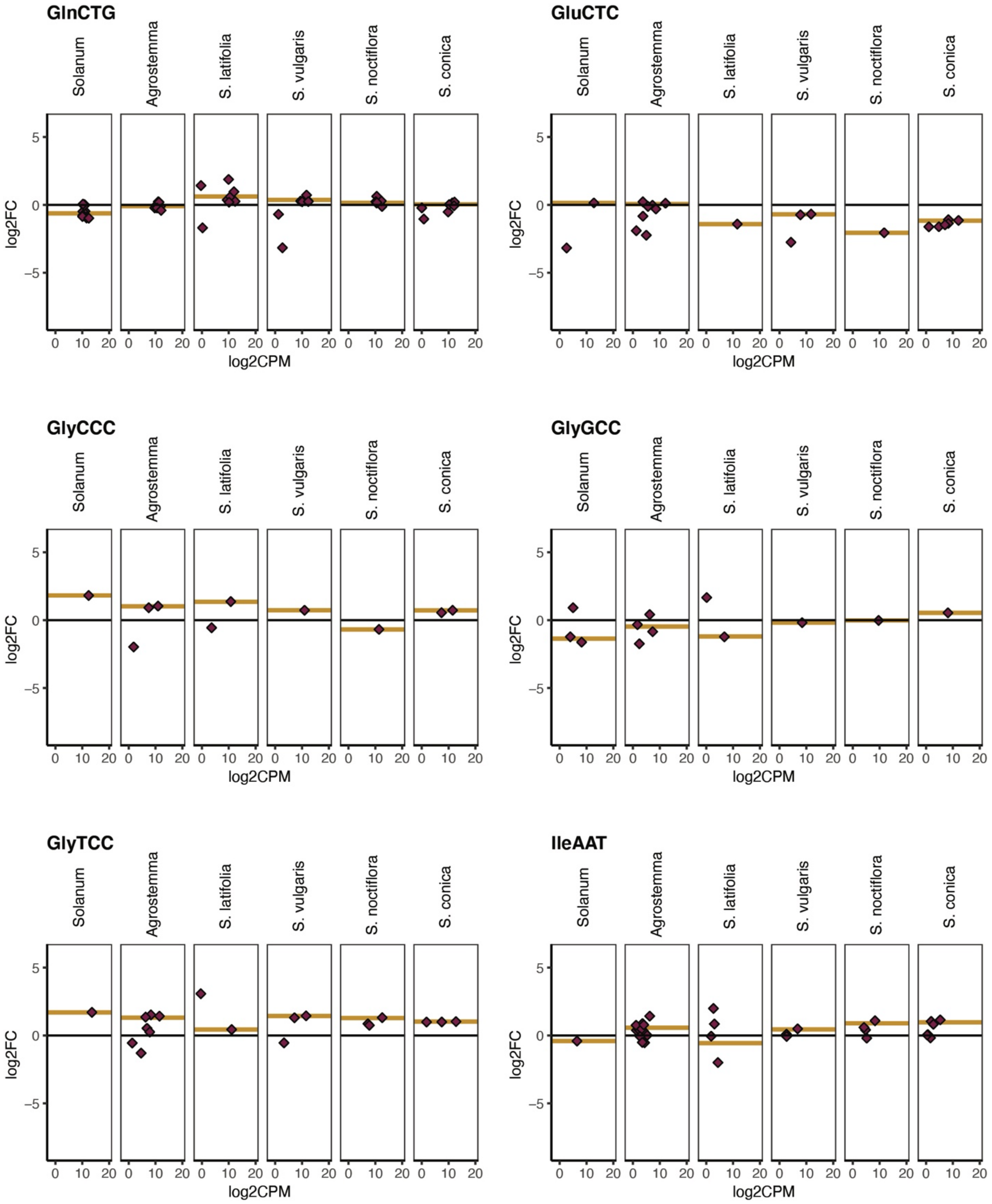
GlnCTG
GluCTC
GlyCCC
GlyGCC
GlyTCC
IleAAT
Solanum
Agrostemma
S. latifolia
S. vulgaris
S. noctiflora
S. conica
log2FC
log2CPM
5
0
−5
0
10
20

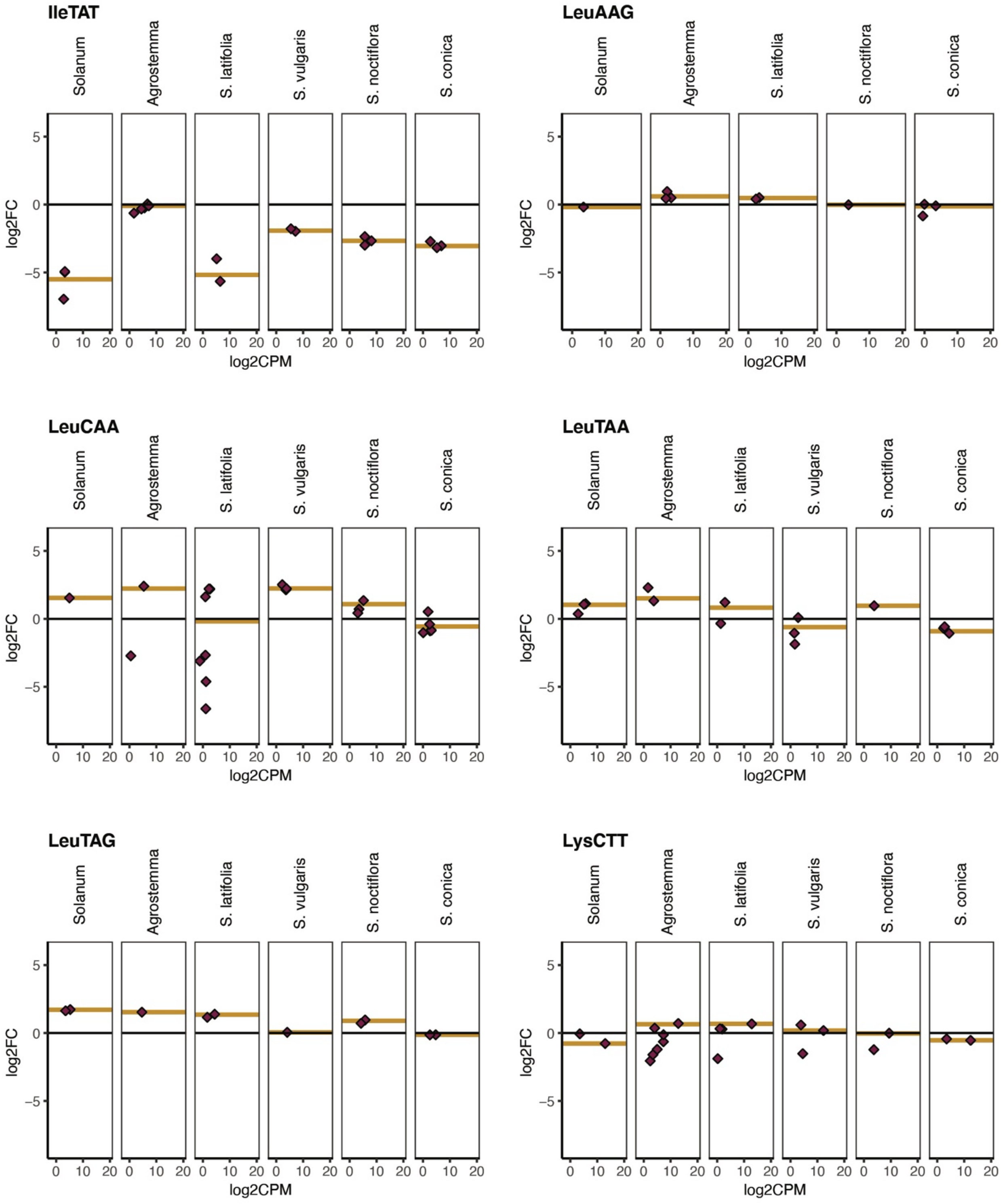
IleTAT
LeuAAG
LeuCAA
LeuTAA
LeuTAG
LysCTT
Solanum
Agrostemma
S. latifolia
S. vulgaris
S. noctiflora
S. conica
log2FC
log2CPM
5
0
−5
10
20

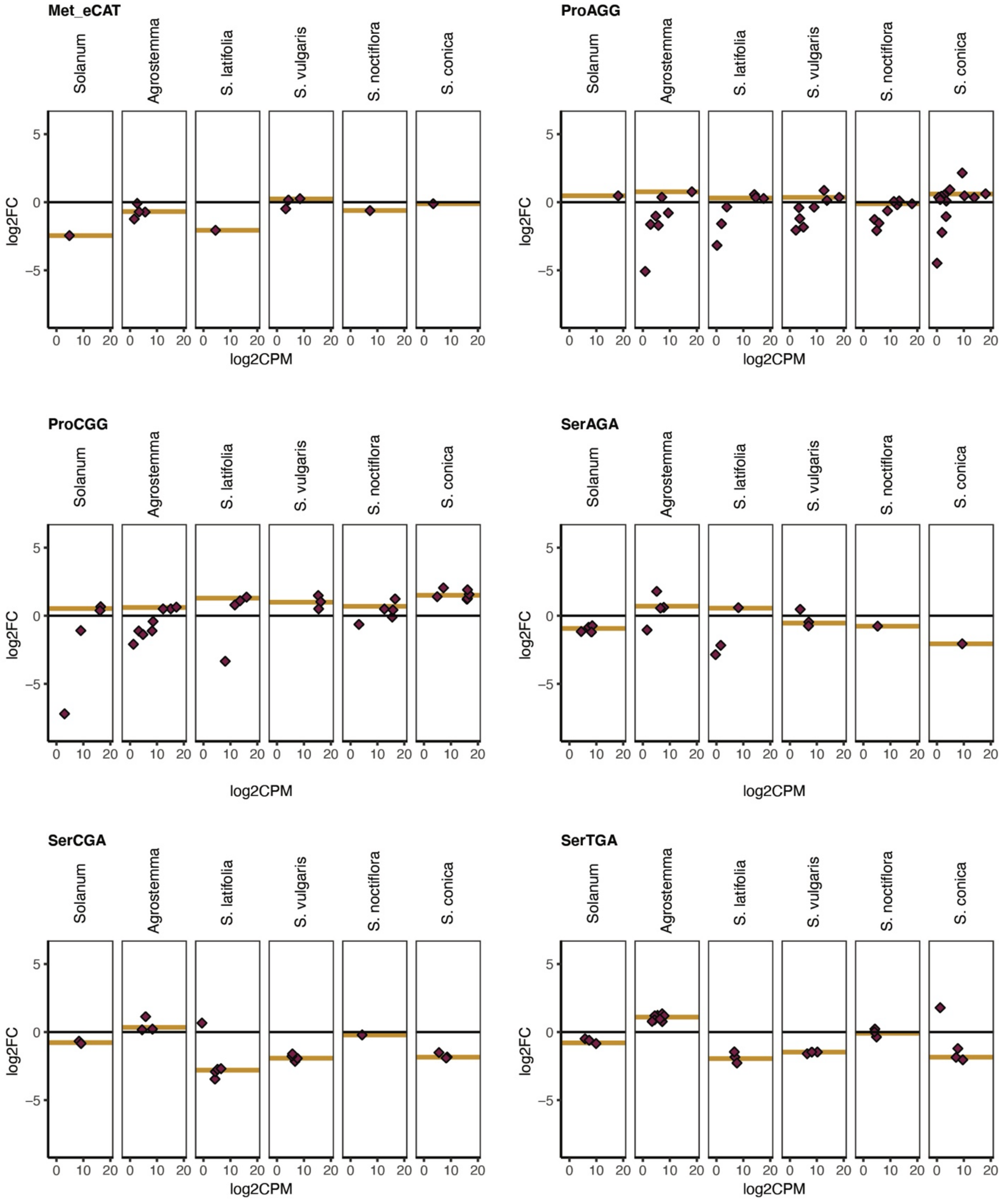
Met_eCAT
ProAGG
ProCGG
SerAGA
SerCGA
SerTGA
Solanum
Agrostemma
S. latifolia
S. vulgaris
S. noctiflora
S. conica
log2FC
log2CPM
5
0
−5
0 10 20

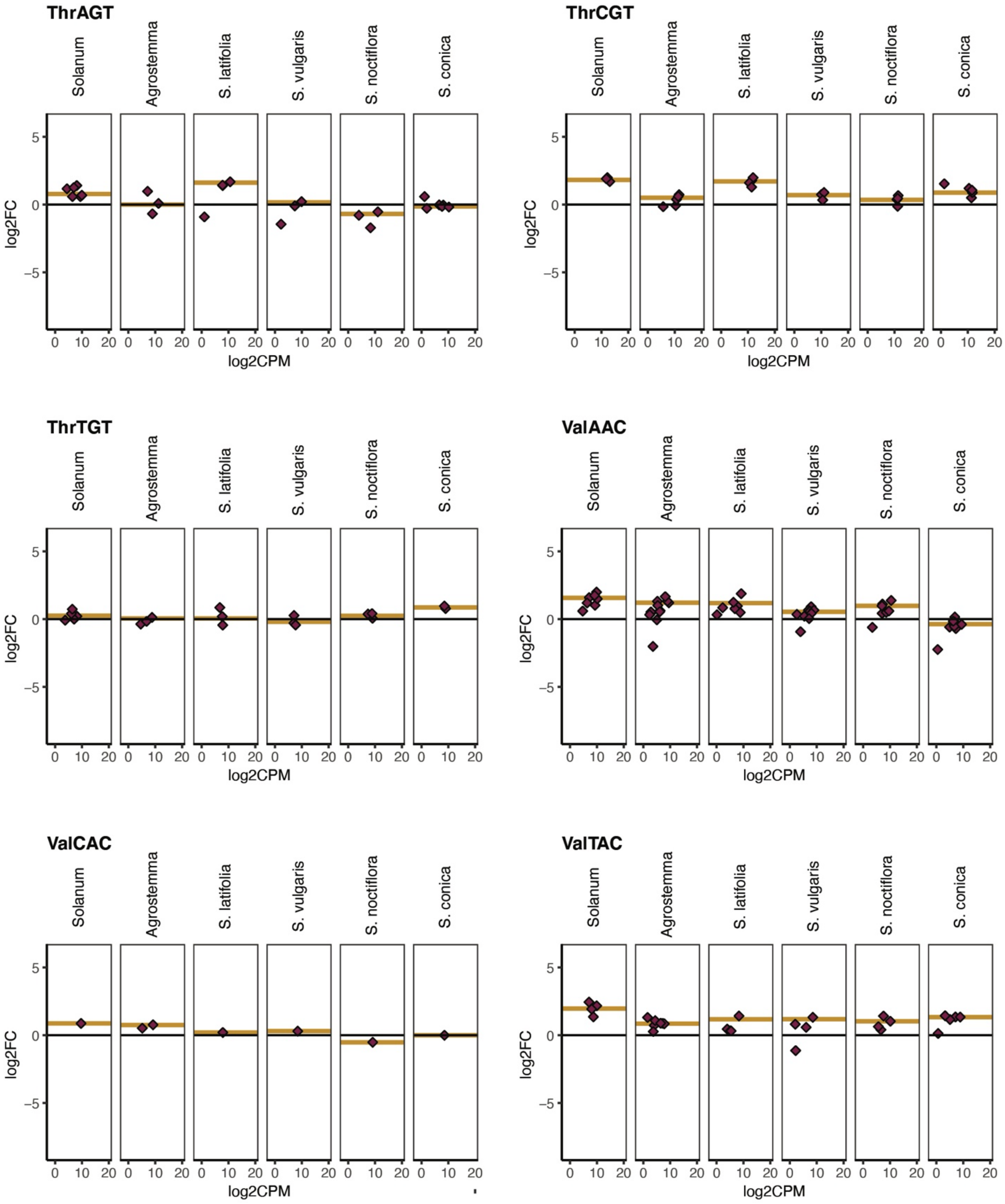


Supp. Fig. 5| Enrichment and expression of individual cytosolic tRNA genes The y-axis is enrichment in $\log_2$ fold change in mitochondrial isolates versus total-cellular samples. The x-axis is expression level in counts per million on a $\log_2$ scale. Points represent individual reference sequences, and gold lines represent the average enrichment of all nuclear-encoded tRNAs with that anticodon weighted by expression (see Methods).
